# Individual connectivity-based parcellations reflect functional properties of human auditory cortex

**DOI:** 10.1101/2024.01.20.576475

**Authors:** M. Hakonen, L. Dahmani, K. Lankinen, J. Ren, J. Barbaro, A. Blazejewska, W. Cui, P. Kotlarz, M. Li, J. R. Polimeni, T. Turpin, I Uluç, D. Wang, H. Liu, J. Ahveninen

## Abstract

Neuroimaging studies of the functional organization of human auditory cortex have focused on group-level analyses to identify tendencies that represent the typical brain. Here, we mapped auditory areas of the human superior temporal cortex (STC) in 30 participants by combining functional network analysis and 1-mm isotropic resolution 7T functional magnetic resonance imaging (fMRI). Two resting-state fMRI sessions, and one or two auditory and audiovisual speech localizer sessions, were collected on 3–4 separate days. We generated a set of functional network-based parcellations from these data. Solutions with 4, 6, and 11 networks were selected for closer examination based on local maxima of Dice and Silhouette values. The resulting parcellation of auditory cortices showed high intraindividual reproducibility both between resting state sessions (Dice coefficient: 69–78%) and between resting state and task sessions (Dice coefficient: 62–73%). This demonstrates that auditory areas in STC can be reliably segmented into functional subareas. The interindividual variability was significantly larger than intraindividual variability (Dice coefficient: 57%–68%, p<0.001), indicating that the parcellations also captured meaningful interindividual variability. The individual-specific parcellations yielded the highest alignment with task response topographies, suggesting that individual variability in parcellations reflects individual variability in auditory function. Connectional homogeneity within networks was also highest for the individual-specific parcellations. Furthermore, the similarity in the functional parcellations was not explainable by the similarity of macroanatomical properties of auditory cortex. Our findings suggest that individual-level parcellations capture meaningful idiosyncrasies in auditory cortex organization.

## 1 Introduction

Differences in behavior and cognition in individual people have been shown to reflect individual-specific variability of patterns of functional connectivity [1–9]. Individual variability may also be one of the major factors underlying why a widely accepted model of the human auditory cortex (AC), analogous to that found in non-human primates [10–13], is still lacking [14]. Microanatomical and functional neuroimaging studies provide evidence that the human AC can be divided into core, belt, and parabelt areas, as well as into parallel anterior vs. posterior feature pathways, which resemble the subdivisions of non-human primate models [11, 15–19]. However, contrasting interpretations exist on the exact layout of these broader arrangements in humans.

In recent years, the development of ultra-high-resolution functional magnetic resonance imaging (fMRI) at 7 tesla (7T) has allowed a deeper investigation into the tonotopy of the primary AC [20–23] and the superior temporal plane [15, 23–26]. Previous 7T fMRI studies have provided converging evidence that there is a large area on Heschl’s gyrus (HG) that is sensitive to low frequencies and is surrounded posteriorly, antero-medially, and antero-laterally by areas sensitive to high frequencies. Interestingly, an individual-level analysis suggested that the AC contains additional frequency reversals that were not evident in the group-level maps of the previous studies [15, 27, 28].

While the other sensory cortices are relatively similar across individuals, the human AC shows larger interindividual anatomical and functional variability [15, 27, 29, 30], a feature that is typical of associative brain areas. AC also resembles associative areas in its anatomy and function, as it includes integrative connections [31], both early and late ACs have several connections to the prefrontal cortex [32], and AC has shown to integrate information from different senses as a multimodal representation [33]. Auditory processing is also strongly modified by attention [34–37]. Moreover, AC is highly plastic, and its function is shaped by experience (for a review, see [38, 39] , which further underlines its interindividual variability. Indeed, AC seems to be an essential part of the networks of brain regions responsible for higher-order cognitive processes, such as language processing [40], working memory [41–44], auditory perceptual decision-making, learning, and prediction [45]. While cortical parcellations have conventionally been generated by spatial normalization and averaging the data over several individuals to increase the signal-to-noise ratio, this approach may not be appropriate for revealing the fine, unique subregions of the human AC since it blurs potentially meaningful interindividual variation. Studying the functional organization of the AC has also been challenged by the relatively large voxel size of conventional 3T fMRI, which may also blur its fine details.

Parcellation approaches relying on individual-specific functional connectivity provide an interesting opportunity to capture the unique functional organizations of individual brains complementing previous studies that have typically defined the regions of interest based on anatomy or population-averaged functional data. Individual-level functional mapping can also enhance group-level analyses since it allows aligning data across participants according to their functional characteristics instead of anatomical landmarks [46]. Indeed, individual-level parcellations have become a very active topic of investigation and a variety of functional connectivity-based approaches have been used to uncover individual differences in the functional organization of the human cortex [47–53]. However, most of the human AC parcellations are still derived from functional specifications (e.g., tonotopic mapping) and microanatomical architecture (e.g., cyto-, myelo-, or receptor architecture) (for a review, see [15]. A recent study investigated subdivisions of Heschl’s gyrus (HG) in individuals using a parcellation approach that relied on structural connectivity derived from diffusion MRI tractography [54]. However, detailed functional connectivity-based parcellations in individuals are still needed.

Here, we investigated functional subareas of the human auditory cortex in superior temporal cortex (STC) in individuals using 1-mm resolution 7T fMRI and a cortical parcellation method that is based on functional connectivity [51] (Fig. 1). The parcellation method identifies functional networks in individuals by iteratively adjusting a population-based functional atlas to fit each individual’s idiosyncratic features. This method has shown to be able to generate highly reproducible parcellations of the whole cortex within individuals and capture meaningful interindividual differences [51]. To validate the test-retest reliability of the results in this study, we measured fMRI data from 30 participants during two resting-state sessions and 1–2 auditory and audiovisual stimulation sessions. Each session was conducted on a different day. We hypothesized that STC can be divided into functional subareas that show high intraindividual reproducibility between sessions, and that the topography and functional specialization of these subareas reveal substantial interindividual variability.

**Figure 1.**
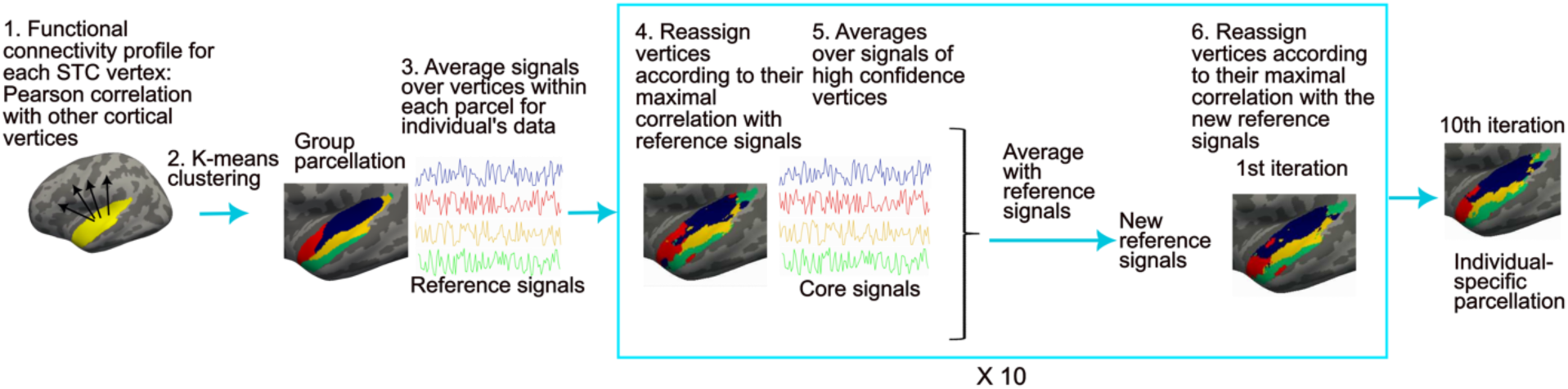
Individual-level functional parcellation method. 1) Pearson correlation was computed between the fMRI signal at each STC vertex and signals of all other cortical vertices in each participant. The individual-level connectivity profiles were averaged across participants at each vertex using a cortical surface-based atlas. 2) Group-level parcellation was computed by using K-means clustering, to cluster vertices with similar connectivity profiles into same network. 3) The individual participant’s fMRI signal time courses were then averaged across the vertices within each group-level network (reference signals). 4) The parcellation was adapted for each individual participant by reassigning vertices to networks according to their maximal correlation with the reference signals. For each vertex, a confidence value was computed as the ratio of the largest and second largest correlations. 5) For each new network of the adapted parcellation, an average was computed using the connectivity profiles of the vertices with confidence values larger than 1.3. The resulting “high-confidence signals” were averaged with the reference signals. 6) Using the resulting new reference signals, the STC vertices were further reassigned to one of the networks. Steps 4–6 were repeated ten times to generate an individual-specific parcellation.

## 2 Results

### 2.1 Auditory cortex parcellations are reproducible within but highly variable across participants

We first generated 23 group parcellations based on functional connectivity by varying the number of clusters in K-means clustering from 2 to 24 (Fig 1, steps 1–2). From these STC parcellations, 4-, 6-, and 11-network parcellations were selected for further analysis based on their group-level Silhouette and Dice values (see Methods; Fig. 2 and Fig. S1 in Supplementary Material). To evaluate the test-retest reliability and intrasubject variability of the parcellations, we generated individual-level parcellations from each of the two resting-state sessions for each participant. For each session, the three STC parcellations were generated (containing 4, 6, and 11 networks). Thereafter, we calculated the overlap between parcellations generated from the two resting-state sessions, for each given cluster solution, by computing their Dice coefficient.

**Figure 2.**
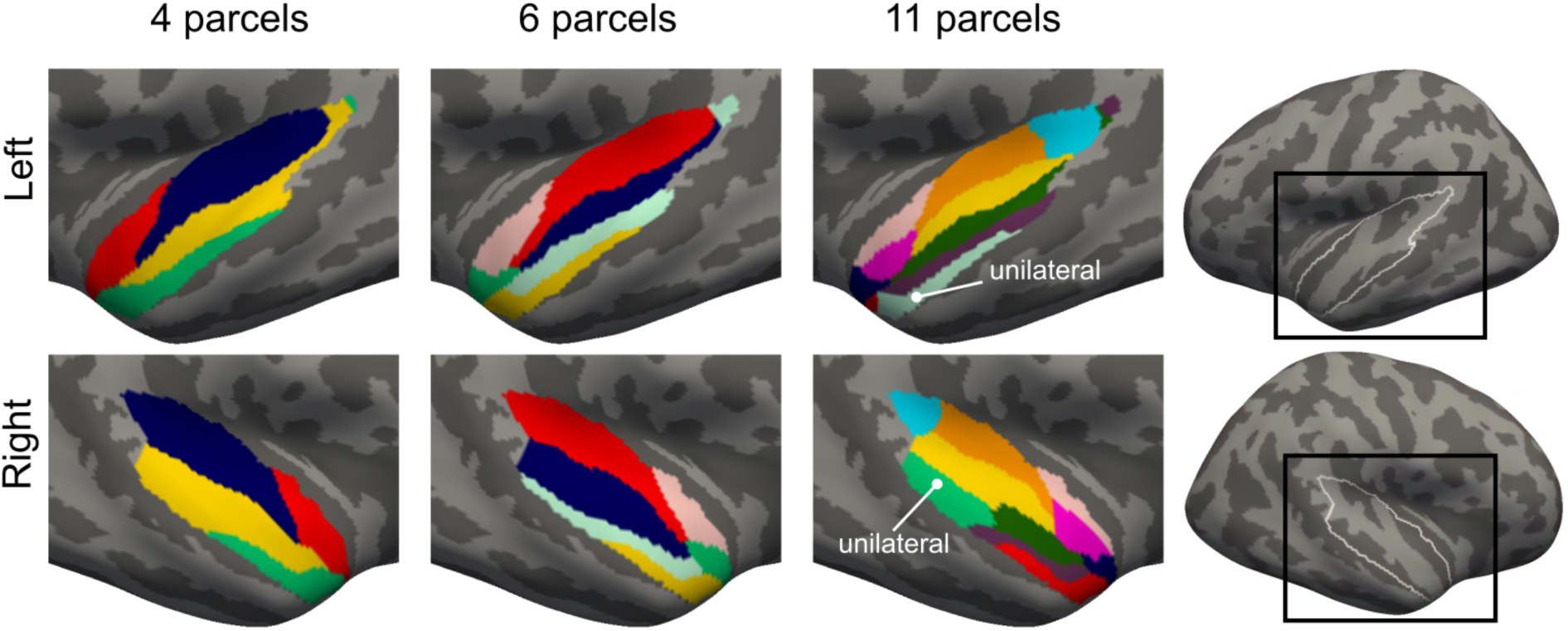
Group-level parcellations of auditory cortices in STC generated using an iterative functional network parcellation method. STC parcellations of 2–24 networks were investigated and three (4-, 6-, and 11-network parcellations) were selected for closer investigation since they yielded local maxima of Silhouette and Dice values at the group level. Different networks are marked with different colors. In the 11-network parcellation 9 of the networks were bilateral and two unilateral.

The parcellations were consistent within individuals between resting-state sessions (Figures 3 and 4, Table S1). At the same time, the parcellations captured high interindividual variability within resting-state sessions. Notably, the interindividual variability was significantly higher than intraindividual variability (*p*<0.001 for each parcellation, the Wilcoxon signed-rank test, corrected for multiple comparisons), indicating that the parcellations were robust and the interindividual variability was related to the actual functional organization of the brain rather than noise. Interindividual variability was significantly higher in the left than in the right hemisphere (4 networks: left: 62±0.8%, right: 68±0.6%; 6 networks: left: 57±0.6%, right: 64±0.6%; 11 networks: left: 57±0.5%, right: 62±0.6%, p<0.001 for all parcellations). Each of the individual networks of the 4-, 6-, and 11-network STC parcellations showed high reproducibility within individuals (Tables S4–S6, Supplementary Material). Again, each and every network was more consistent within than between individuals (Table S4, Supplementary Material), indicating that each network has substantial interindividual variability.

**Figure 3.**
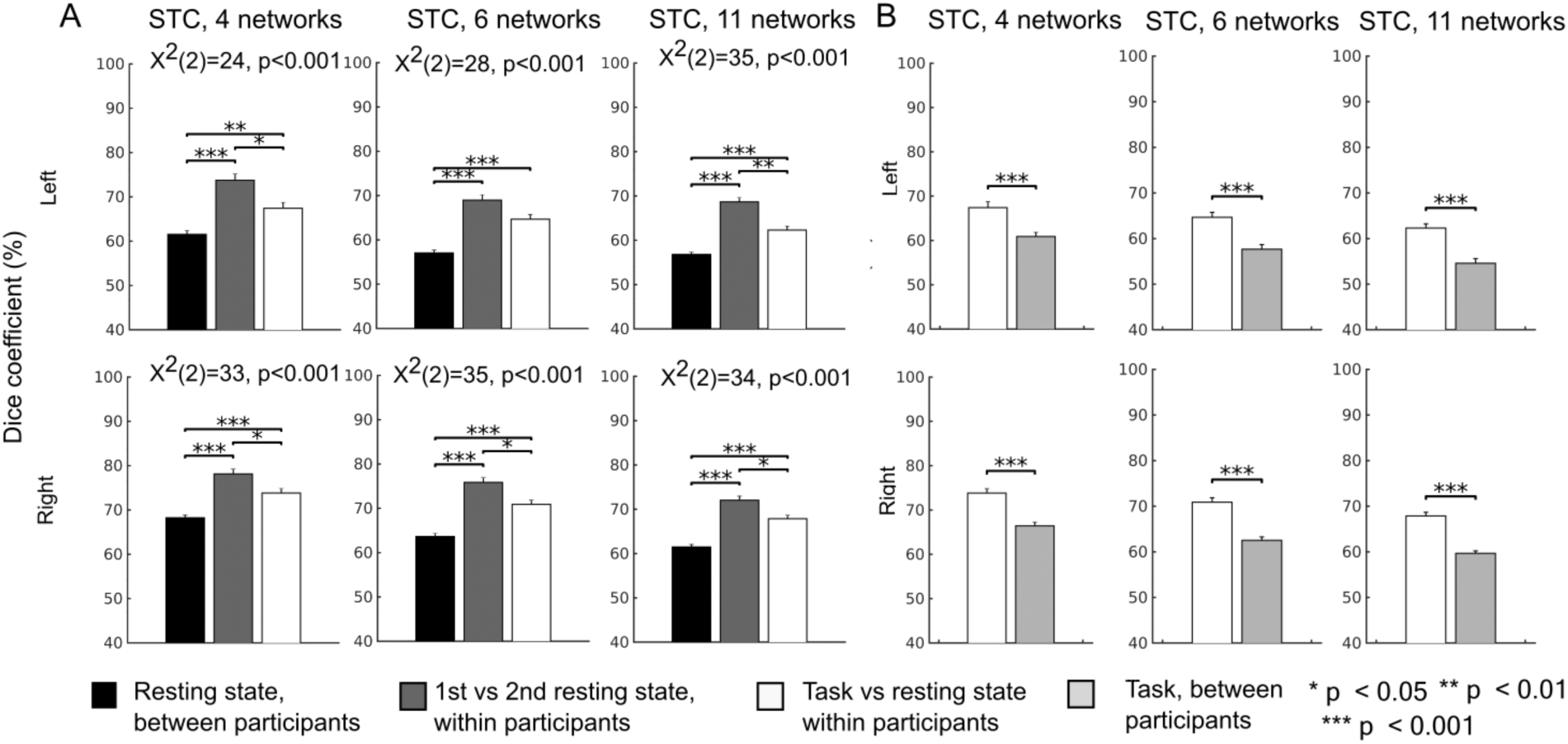
Intraindividual and interindividual variability of the parcellations. A) The parcellations generated using the resting-state sessions demonstrated high intraindividual reliability and high interindividual variability. When comparing two resting-state sessions of the same participant, on average, 69–78% of the STC vertices were assigned to the same networks. Between a given participant and any other participant, on average, only 57–68% of the STC vertices were assigned to the same networks. For each STC parcellation, the consistency of network membership was significantly higher within than between participants (p<0.001 for all conditions). The overlap between the resting-state parcellations within participants was higher than that between the resting-state and task parcellations (task vs. rest: 62–74%, rest vs. rest: 69–78%, p<0.05) for all parcellations expect the left hemispheric 6-network parcellation. This suggests that auditory cortex functional networks can be shaped by task. B) The correspondence between task and resting-state parcellations of a given participant (62–74%) was higher than the correspondence between task parcellations of the participant and any other participant (55–66%; p<0.001), suggesting than the parcellations are most similar within participants regardless of the task. The differences between these three conditions were estimated with the Friedman test and pairwise Wilcoxon signed rank tests. The results were corrected for multiple comparisons using Benjamini-Hochberg procedure [55]. Error bars indicate standard errors of the mean (SEM).

**Figure 4.**
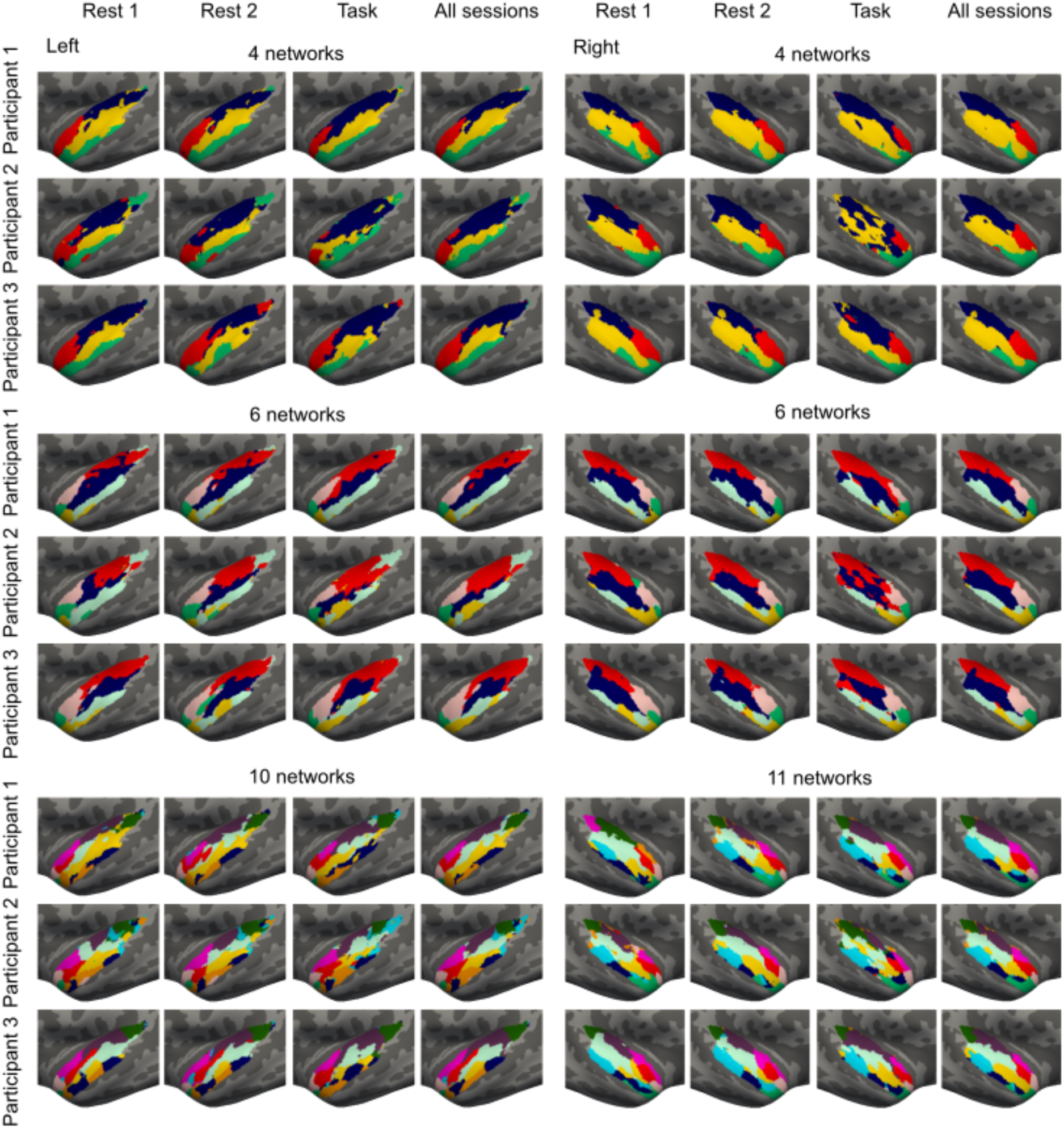
Individual-level 4-, 6-, and 11-network STC parcellations for three representative participants. The parcellations were generated separately for two resting-state fMRI sessions and for the session during which the participant was performing auditory and audiovisual tasks (see Methods for task descriptions). All sessions (i.e., Rest 1, Rest 2, and Task) were measured on different days. In addition, we generated parcellations using data from all three sessions.

To test whether the parcellation algorithm also captures variability in the cortical thickness and curvature, we conducted Mantel test between the dice values and between-participant similarities of the STC thickness and curvature values. The similarity in the anatomical measures was estimated with Pearson correlation of the thickness and curvature values between participants, resulting one similarity matrix for thickness and one for curvature (p=0.34-0.83). The dice coefficients were computed using the parcellations created based on all rest and task sessions. Similarity in the parcellations did not correlate with the similarity of the thickness or curvature, suggesting that the interindividual variation mainly reflects functional rather than anatomical variability.

### 2.2 Auditory cortex parcellations are most consistent within individuals regardless of the task

To investigate the reproducibility of the parcellations between different functional tasks, we compared the resting-state parcellations with the parcellations derived from the fMRI data measured during auditory and audiovisual speech processing tasks. STC parcellations were relatively consistent between resting-state and task sessions within participants (Fig. 3B, Table S2). Critically, the task parcellation overlapped more precisely with the resting-state parcellation of the same participant than with the task parcellations of the other participants (p<0.001, Fig. 3B, Table S2): this suggests that the parcellations are most similar within individuals, independently of whether or not the participant was performing a task during scanning (Fig. 3B, Table S2). The overlap between task and resting-state parcellations was also higher within participant than the overlap of the resting-state parcellations between participants in all cases except in the right-hemispheric 4-network parcellation and the left hemispheric 6-parcel parcellation (p<0.001, Fig. 3A). However, within participants, the two resting-state parcellations overlapped more precisely than resting-state and task parcellations (Fig. 3A, Table S3, p<0.05) for all expect the left-hemispheric 6-network parcellation. This suggests that task can modulate functional networks of STC.

### 2.3 Task activations are more homogeneous within individual-specific than group parcellations

fMRI data were also collected during basic auditory stimulation and audiovisual speech processing to compare functional parcellations to task-based data. Tonotopy and amplitude modulation rate representation were mapped orthogonally using the data from a task where the participants were presented with serial blocks of sounds, which varied both in their center frequency (tonotopy mapping) and in their amplitude modulation rate (rate representation mapping). In Figures 5 and 6, “Auditory vs. baseline” is the contrast between all auditory stimuli and no auditory stimulation, “High vs. low frequency” between high (1.87 kHz or 7.47 kHz) and low (0.12 kHz or 0.47 kHz) carrier frequencies, and “Slow vs. fast AM” between slow (4 cycles/s) and fast (32 cycles/s) amplitude modulations. The results for slow and fast AM rates vs. baseline are shown in the Supplementary Material (Fig. S2). To localize the cortical areas responding to speech and audiovisual interaction, the participants were presented with clear or blurred video clips of a person voicing either “rain” or “rock.” The auditory component in the videos was either acoustically intact or replaced with noise that matched the spectrotemporal power distribution of the original speech. The “Speech vs. noise” contrast in Figures 5 and 6 refers to the contrast between all conditions with clear speech and all with noise. “Visual input enhances noisy speech processing” refers to the areas where response to noisy speech was stronger when visual input was clear.

**Figure 5.**
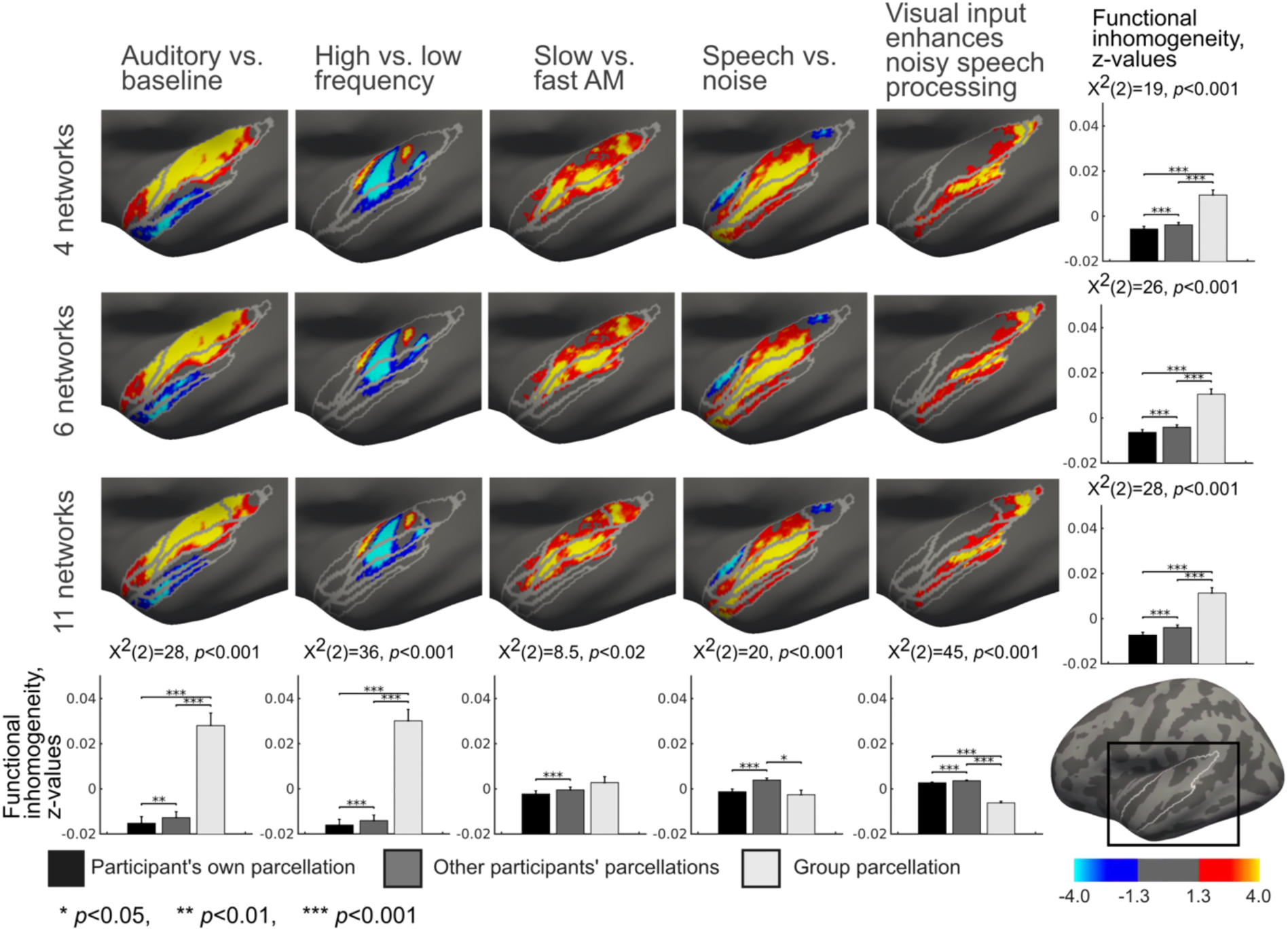
Topographic correspondence between the STC parcellations and GLM task contrasts in the left hemisphere. The group-level parcellation is overlaid on the group-level GLM contrasts computed from auditory and audiovisual tasks. The bar diagrams in the right column show functional inhomogeneity averaged over all contrasts within each of the three parcellations. The bar diagrams in the bottom row show functional inhomogeneity averaged over all parcellations within each of the five contrasts. The functional inhomogeneity was computed between the individual GLM contrast map of each participant and 1) their own individual–specific parcellation, 2) individual-specific parcellation of all other participants, and 3) group-average parcellation. The differences between these three conditions were estimated with the Friedman test and pairwise Wilcoxon signed rank tests. The results were corrected for multiple comparisons using Benjamini-Hochberg procedure [55]. Error bars indicate standard errors of the mean (SEM).

**Figure 6.**
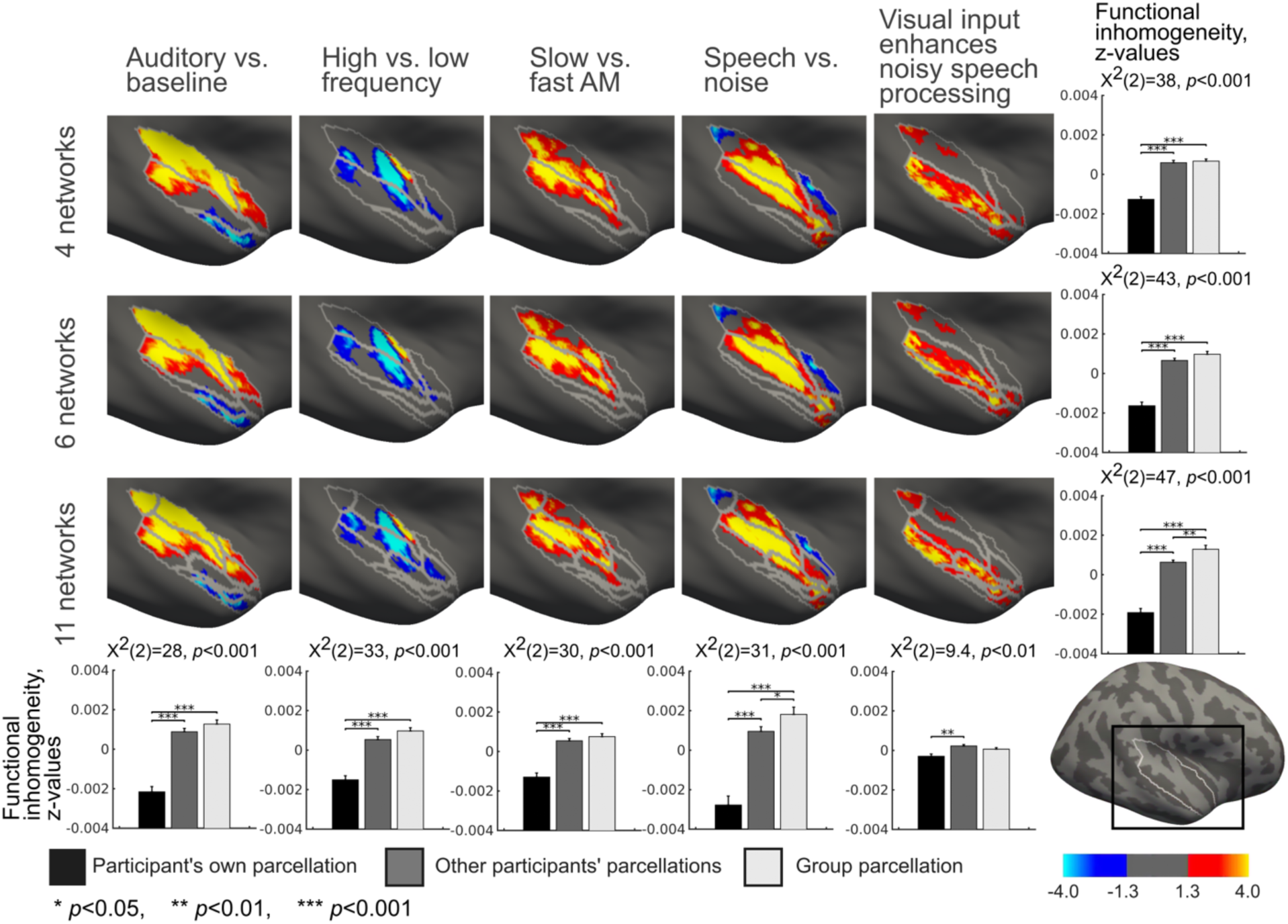
Topographic correspondence between the STC parcellations and GLM task contrasts in the right hemisphere. The group-level parcellation is overlaid on the group-level GLM contrasts computed from the auditory and audiovisual tasks. The bar diagrams in the right column show functional inhomogeneity averaged over all contrasts within each of the three parcellations. The bar diagrams in the bottom row show functional inhomogeneity averaged over all parcellations within each of the five contrasts. The functional inhomogeneity was computed between the individual GLM contrast map of each participant and 1) their own individual-specific parcellation, 2) individual-specific parcellation of all other participants, and 3) group-average parcellation. The differences between these three conditions were estimated with the Friedman test and pairwise Wilcoxon signed rank tests. The results were corrected for multiple comparisons using Benjamini-Hochberg procedure [55]. Error bars indicate standard error of the mean (SEM).

If the parcellations represent functional areas, task response patterns can be expected to align with parcellation boundaries. Figures 5 and 6 show the network parcellations overlaid with auditory and audiovisual speech fMRI task data. Based on visual inspection, the parcellation boundaries are reasonably well aligned with task response patterns. To quantitatively estimate the topographic correspondence between parcellations and task response patterns, we computed functional inhomogeneity for each parcellation. Functional inhomogeneity was defined as an average over network-specific standard deviations. The network-specific standard deviations were computed as the standard deviation of the task responses (*z*-scores) within each network, adjusted with the network size (see Methods). Low functional inhomogeneity indicates that parcellation aligns accurately with task response patterns. The functional inhomogeneity was estimated between each participant’s task response map and 1) individual-specific parcellation of the same participant, 2) individual-specific parcellations of the other participants, and 3) group-averaged parcellation.

When averaged across five task contrasts, the individual-specific parcellation was functionally more homogeneous than other participants’ parcellations or group parcellation for all parcellations (*p*<0.001, Fig. 5 and 6). This demonstrates the highest correspondence between individual-specific parcellation boundaries and individual task response patterns. When the functional inhomogeneities were averaged across the STC parcellations within contrasts, the individual-specific parcellation was functionally the most homogeneous (p<0.01, Fig. 5 and 6, Fig S2) for all except three contrasts in the left (Auditory vs. baseline, Slow vs. fast AM, Speech vs. noise, Visual input enhances noisy speech processing) and one in the right hemisphere (Visual input enhances noisy speech processing). For “Slow vs. fast AM” contrast in the left and “Visual input enhances noisy speech processing” contrast in the right hemisphere the participant’s own parcellation was functionally more homogeneous than other participants’ parcellations, but no statistical differences were found between these parcellations and the group level parcellation. For “Speech vs. noise” contrast in the left hemisphere, no differences were found between participant’s own and group parcellations but these both were functionally more homogeneous than other participants’ parcellations. For “Visual input enhances noisy speech processing” contrast in the left hemisphere, the group parcellation was functionally most homogeneous and the participant’s own parcellation more homogeneous than other participants’ parcellations.

To further investigate the functional specificity of auditory areas in STC, we calculated the percentage of the total contrast effect size that each network of each parcellation explained. Figure 7 shows that the task contrast effects were mostly restricted within certain networks, suggesting specificity of the STC networks. The correlation of the network-specific percentages of the total contrast effect size between task contrasts was lowest for the 11-network parcellation (left: 4 networks: 0.86, 6 networks: 0.84, 11-networks: 0.62; right: 4 networks: 0.90, 6 networks: 0.93, 11-networks: 0.83). This suggests that functional specificity was highest for the 11-network parcellation.

**Figure 7.**
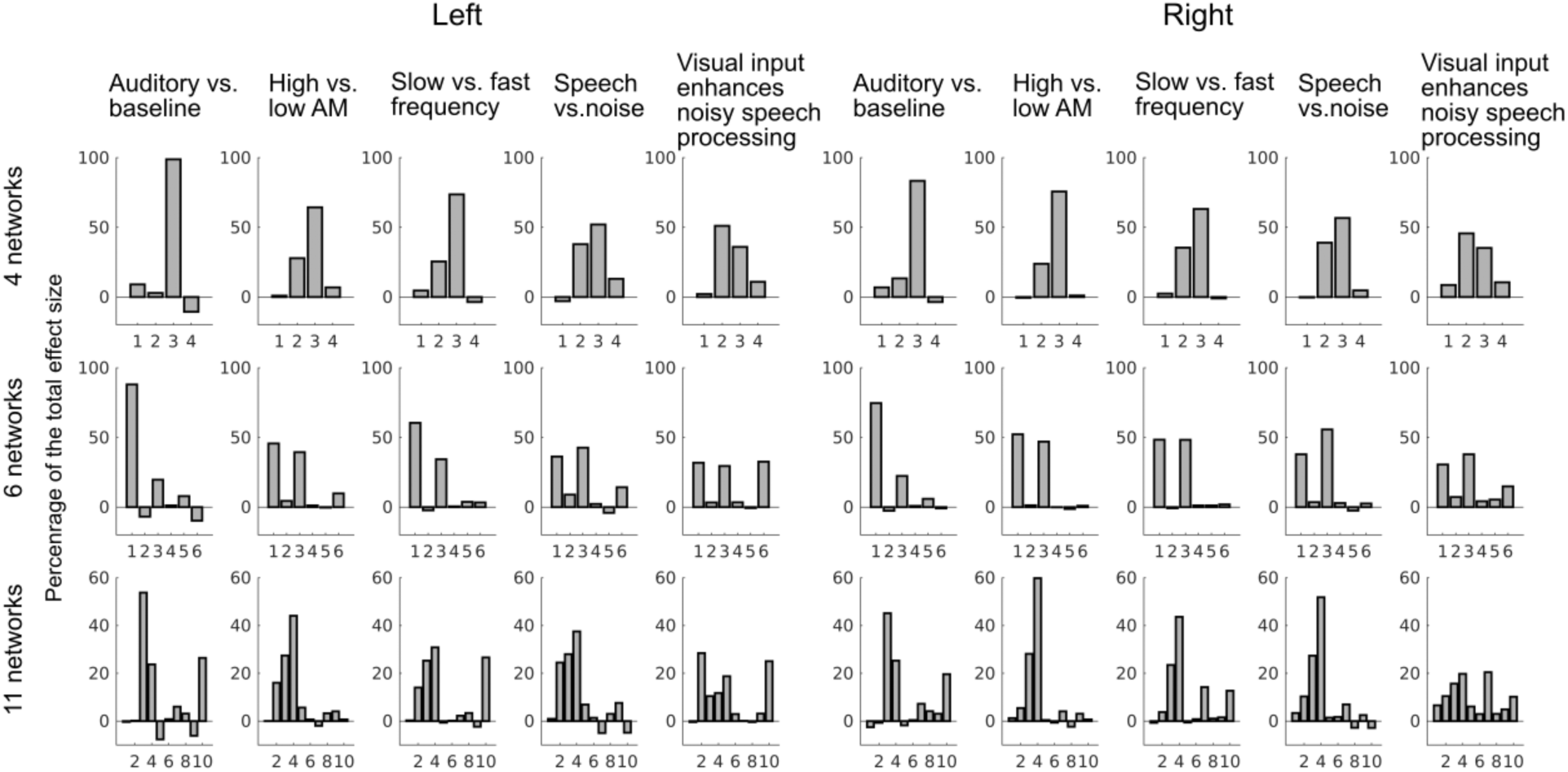
Functional specificity of the STC parcellations. The percentage of the total effect size is shown for each network in each parcellation. The task effect was concentrated within certain networks, suggesting functional specificity of the parcellations.

### 2.4 Connectional homogeneity is higher within individual-specific than group-parcellations

To further evaluate the neurophysiological plausibility and validity of the parcellations, connectional homogeneity was estimated using individual resting-state data. To estimate connectional homogeneity, Pearson correlation coefficients were calculated between all pairs of vertices within each network and transformed into z-values using Fisher z-transform. The resulting z-values were averaged within each network and, thereafter, an average was computed across network-wise z-values while accounting for the network size (see Methods). For each participant, the connectional homogeneity was estimated using their resting state fMRI and 1) their individual-specific parcellation, 2) the individual-specific parcellations of the other participants, and 3) the group-average parcellation. In this analysis we used parcellations generated based on data from all three fMRI sessions. The connectional homogeneity was higher for the individual-specific STC parcellations than for other participants’ parcellations or for the group parcellation (Fig. 8, *p*<0.001 for all parcellations), demonstrating that the individual-specific parcellations have the highest plausibility and validity. Further, the other participants’ parcellations yielded higher connectional homogeneity than the group-average parcellation (*p*<0.01 for all parcellations) for all expect the right-hemispheric 6-network parcellation for which no differences were found between these two parcellations.

**Figure 8.**
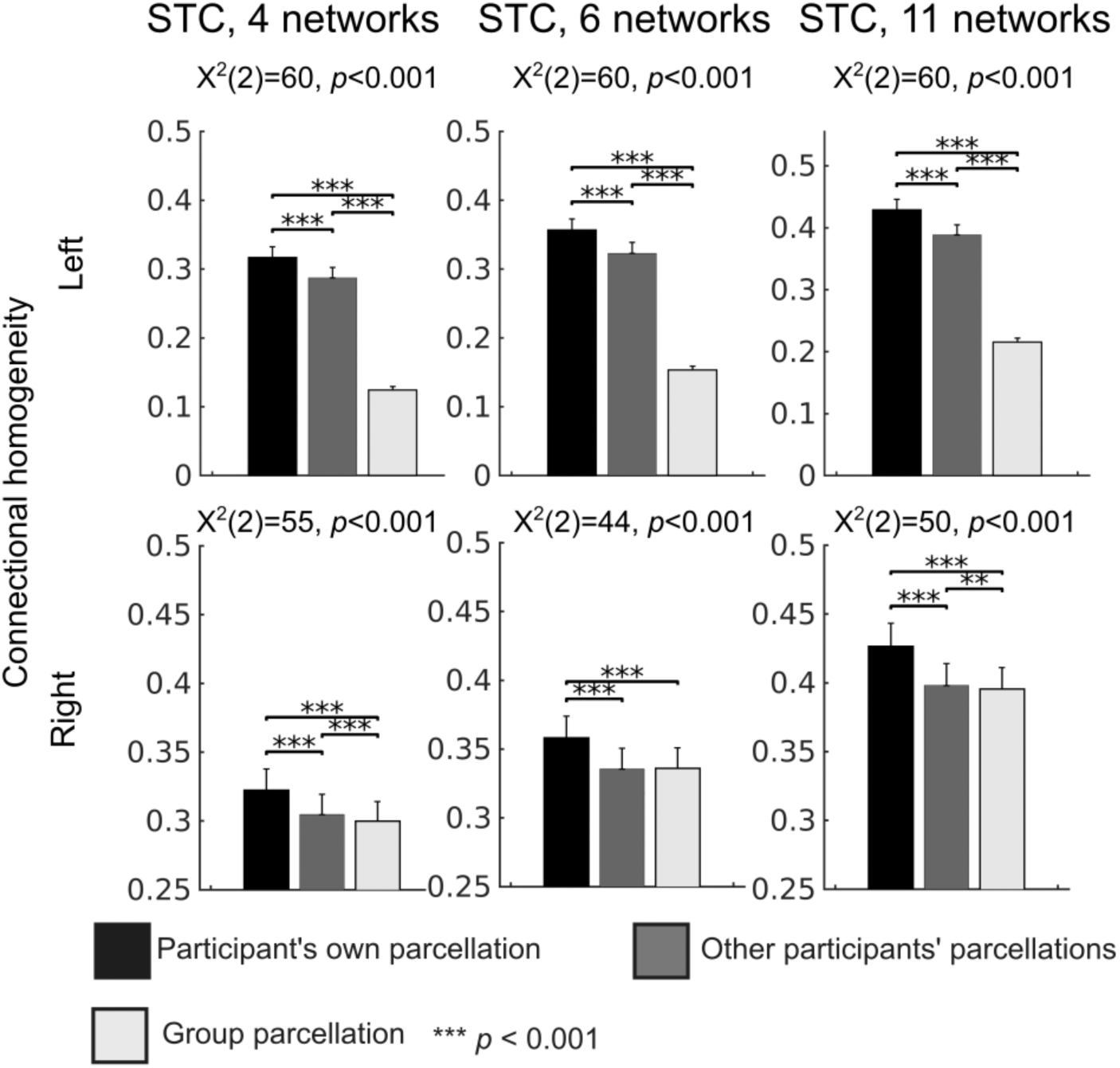
Connectional homogeneity of the STC parcellations. For each individual, connectional homogeneity was estimated using 1) their individual-specific parcellation, 2) the individual-specific parcellations of the other participants, and 3) the group-average parcellation. The statistical differences were estimated with the Friedman test and pairwise Wilcoxon signed rank tests. The results were corrected for multiple comparisons with Benjamini-Hochberg procedure [55]. The error bars indicate standard errors of the mean (SEM).

### 2.5 Seed-based functional connectivity maps are specific to auditory cortex networks

The parcellation method classifies vertices with similar connectivity profiles into the same network. We conducted seed-based connectivity analysis using each network in turn as a seed region (an average signal was computed over the network vertices) to determine the cortical areas with which the STC networks were connected (Fig. 9, Supplementary Material: Fig. S3 and Fig. S4). As can be expected, the networks were connected with traditional speech and language areas including temporal cortices, Broca’s area, and dorsal precentral areas [56, 57]. Some of the networks were also connected with lateral occipital/occipitotemporal cortices, including the human middle temporal area (hMT) that is associated with visuospatial motion processing [58, 59]. The networks were also consistent with previous studies showing a dissociation between anterior/ventral "what" (Fig. 9, Networks 5-7, 10) and posterior/dorsal "where" pathways (Networks 2, 11) [17, 19, 60]. Moreover, the connectivity profiles of the adjacent STC networks showed statistically significant differences between each other even in the most detailed 11-network parcellation (Fig. 10).

**Figure 9.**
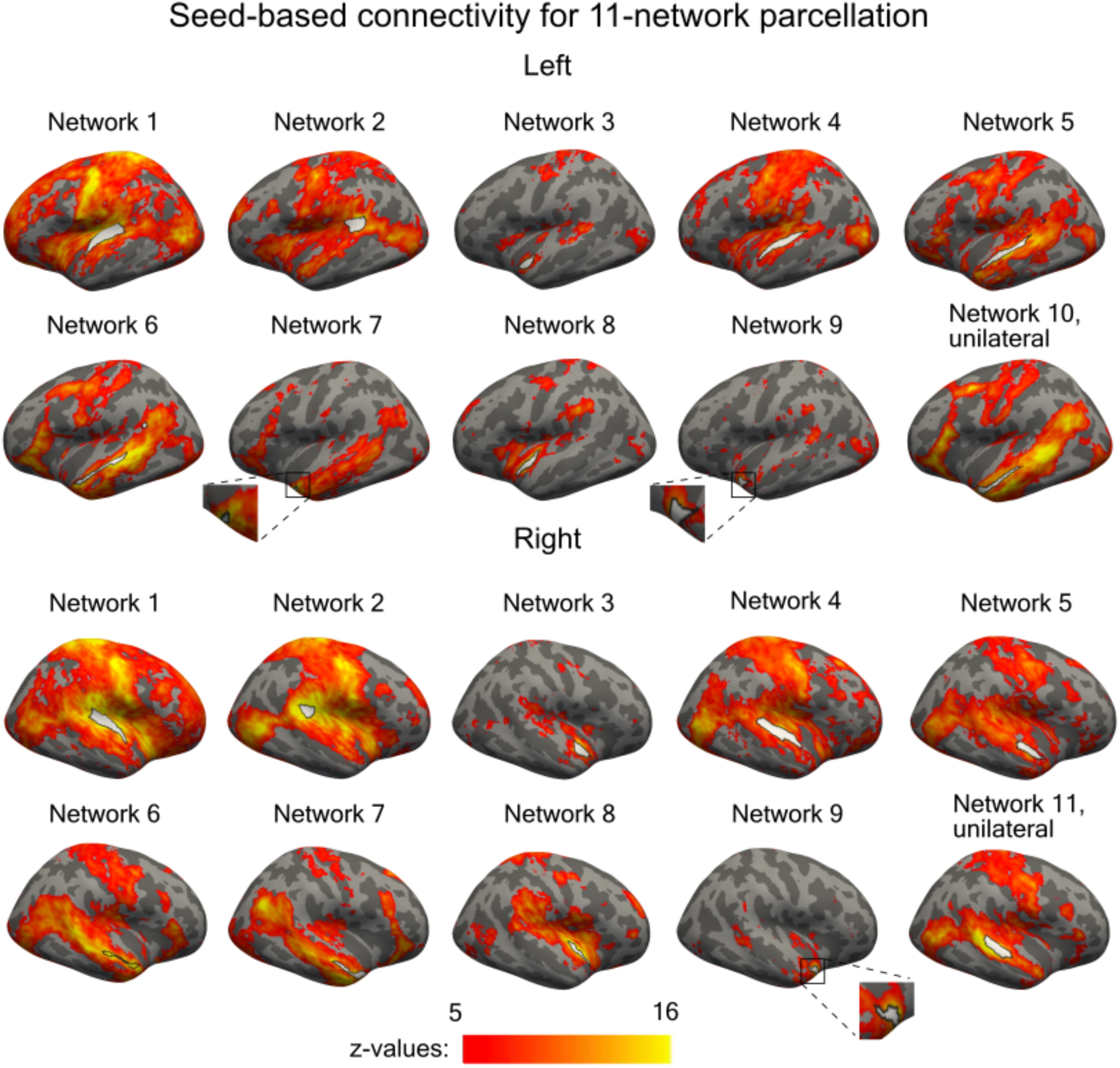
Seed-based functional connectivity maps for each network in the 11-network parcellation. Network 10 was only found in the left and Network 11 in the right hemisphere. Other networks were bilateral. One-sample t-test was performed for each vertex. The presented maps are thresholded at p<0.05 and corrected for multiple comparisons with cluster-extent based permutation thresholding with a cluster-forming threshold of p<0.001 (one-sample T-test).

**Figure 10.**
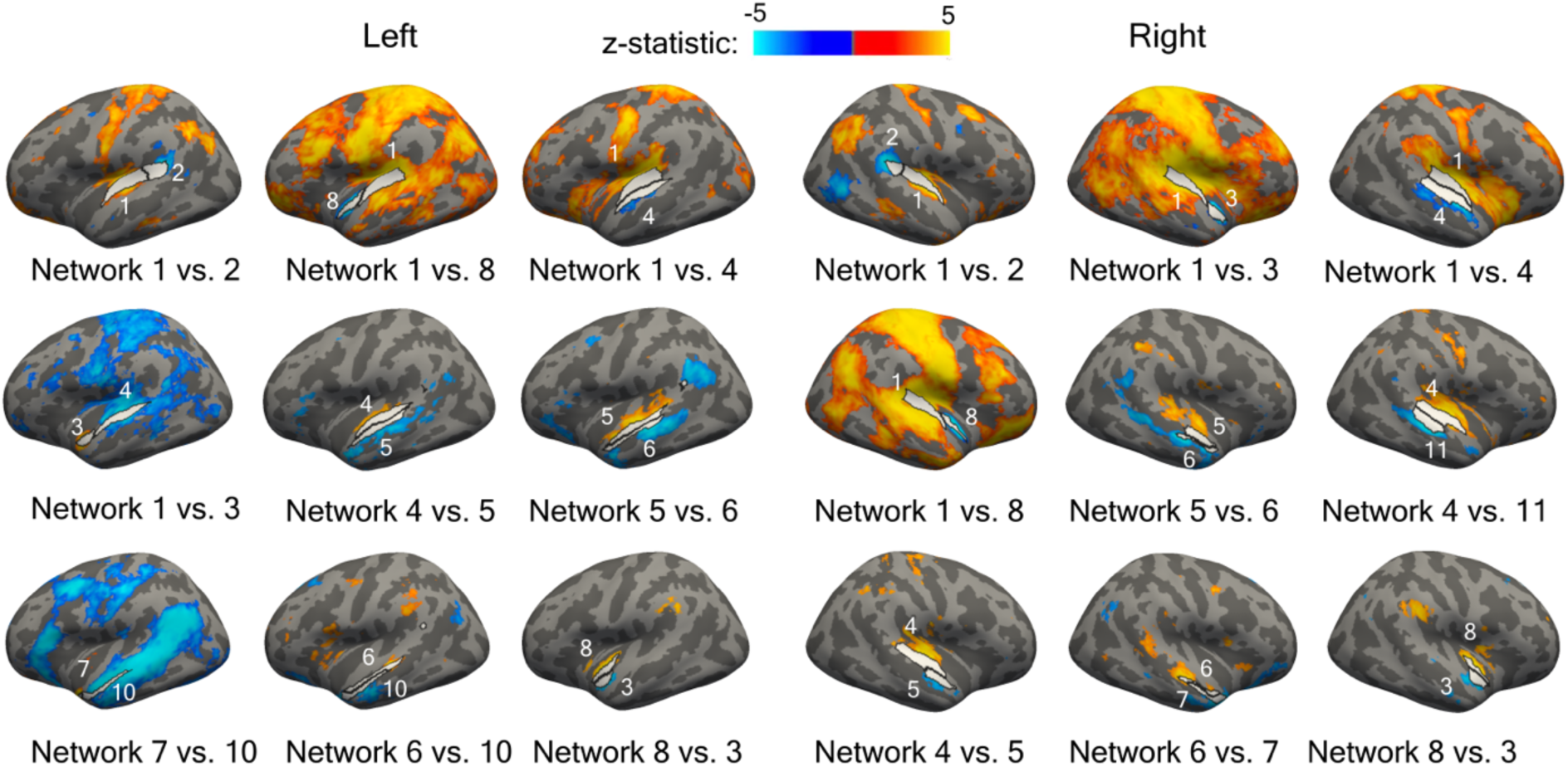
Differences between seed-based functional connectivity maps for pairs of adjacent networks in the 11-network parcellation. The Wilcoxon signed-rank test was calculated vertex wise between the connectivity maps of adjacent network pairs at each vertex. The resulting p-values were adjusted for multiple comparisons using Benjamini-Hochberg method [55]. The threshold was set at p < 0.05.

## 3 Discussion

Our results suggest that human auditory areas in STC can be reliably divided into fine-grained subareas in individuals by using functional network analysis and ultra-high resolution 7T fMRI. The subarea arrangements were highly reproducible within individuals but showed substantial interindividual variability (*n*=30). Moreover, the parcellations were functionally homogeneous and aligned with response patterns elicited by auditory and audiovisual tasks, which is indicative of their neurophysiological plausibility and validity. This is a notable extension to the auditory cortex parcellation literature, which has mainly focused on tonotopic mapping or group-level parcellations derived from anatomical data (for a group-level functional-connectivity based STC parcellation, see [61] for individual-level anatomical parcellation, see [54].

### 3.1 The same auditory cortex subareas can be reliably identified in individuals at multiple network resolutions

Thus far, most existing parcellation of human auditory cortex have been derived from microanatomical architecture or using tonotopic mapping (for a review, see [15]. However, our recent study provided initial evidence that the human STC could be divided into subareas, at the group level, based on cortico-cortical or cortico-cerebellar resting-state functional connectivity [61]. The current study extends previous research by generating individual-level cortico-cortical functional connectivity-based parcellations for STC at different network resolutions.

Another important extension to previous literature is that our study includes task and resting-state fMRI data from the same participants, which allowed us to evaluate the functional properties of the networks. The 4-, 6, and 11-network STC parcellations selected in the individual-level analysis were highly reliable within participants as indicated by a test-retest reliability of 69– 78% for the resting-state data. Moreover, the parcellation boundaries were aligned with GLM topographies elicited by auditory and audiovisual tasks, which further supports their neurophysiological validity. Interestingly, the shape and size of the networks also differed between hemispheres, which is in line with the known hemispheric lateralization of the auditory areas.

Our results may roughly correspond with cytoarchitectonic parcellations determined using human post-mortem brains [62–65]. In the 11-network parcellation (Fig. 2), the turquoise network may correspond to Te2, orange network Te1, light pink TI, and yellow Te3 in the cytoarchitectonic STC parcellation determined recently using histological sections of ten postmortem brains [63]. The dark green, dark violet, and light blue networks may correspond to STS1 and STS2. Ren et al. suggested partly similar correspondence for their group-level functional connectivity-based STC parcellation.

While the 4 and 6-network parcellations were fully bilateral, the 11-network parcellation showed two auditory cortex subareas that were reliably present in only one hemisphere (Fig. 2 and Fig. 9). In the seed-based analysis, the left-hemispheric Network 10 was connected to the classic speech processing networks including Broca’s and dorsal precentral areas, consistent with the known lateralization of human language pathways [56, 57]. The right-hemispheric Network 11 was, in turn, connected to lateral occipital/occipitotemporal cortices, including the human middle temporal area (hMT) that is associated with visuospatial motion processing [58, 59]. Network 11 could, thus, play a role in providing crossmodal visuospatial information to auditory cortex, to support auditory motion processing. The lateralization of this network is consistent with previous studies on auditory motion processing [66].

Based on previous studies on humans and nonhuman primates, STC could include over 10 subareas within a limited region of the superior temporal cortex [11, 13, 29, 67–70]. Tonotopic mapping studies have typically identified three regions at the group level, but results in individuals suggest more fine-grained structure with additional frequency gradients [15]. Ren et al. selected 2- and 6-cluster functional network-based parcellations for STC based on Silhouette values. Further, several metrics and algorithms have been proposed for estimating the number of networks for connectivity-based parcellations [71–74]. However, given the hierarchical multi-resolution organization of the human brain, the optimal number of networks may depend on the brain function of interest. Therefore, this study proposed three STC parcellations with network numbers of 4, 6, and 11. These parcellations showed local maxima of test-retest reliability and cluster separability among 2–23 network solutions, but other solutions may still be valid depending on the application, and the optimal number of networks may vary even between participants.

### 3.2 Parcellations reflect functional specificity of auditory cortex

The parcellation subarea outlines were well aligned with the GLM contrast topographies (Fig. 5 and 6). Many of the task-based contrast effects were largely concentrated within a few parcels only (Fig. 7). The 11-network parcellation appeared to align with the task contrasts most accurately. We also computed the percentage of total task effect size explained by each network. The correlation of these network-wise percentages between tasks was lowest for the 11-network parcellation. This suggests that the task determined more strongly for the 11-network parcellation within which networks the contrast effects were restricted than for the 4- or 6-network parcellations. Overall, the concentration of task-specific effects within selected parcels supports an interpretation that these parcels could reflect functionally relevant auditory subdivisions of STC.

The parcellation followed borders of the auditory processing hierarchy. The cochleotopic areas selective to high or low frequency were bilaterally restricted within the network (marked with orange in Fig. 2), which included medial aspects of STC and early auditory areas of HG. The most significant responses to fast AM vs. baseline were also concentrated in this network. The two adjacent networks in the lateral (yellow) and posterior (turquoise) directions from the earliest areas responded more strongly to slow than fast AM, and also showed a strong contrast to auditory speech vs. noise signals. These results agree with previous evidence that areas lateral to early auditory cortices integrate information over longer timescales and are more sensitive to complex than simple sound attributes [75, 76]. However, in line with [77], no evidence for a clear large-scale amplitude modulation rate topography [78–80] was found.

The area responding more strongly to speech than noise extended to two left-hemispheric networks in the lateral STG and the superior temporal sulcus (STS; dark violet and light turquoise). These networks showed no AM-rate or frequency selectivity, which is consistent with the notion that these areas respond more strongly to words than tones as well as to speech than non-speech sounds [81].

Within the most posterior (turquoise) and lateral (left: dark green, purple; right green, dark green) networks, the contrast between clear vs. noisy visual stimulus was stronger when the accompanying auditory speech signal was noisy than when it was clear. In other words, in these subareas, visual gestures had strongest effect on speech processing when the auditory signal is noisy. This may reflect networks that use multisensory information in challenging listening conditions. This finding is consistent with previous studies using the same stimuli in intracranial recordings in human participants, which reported that posterior areas of non-primary auditory cortex are specifically important for multisensory integration of noisy speech signals [82, 83].

### 3.3 Auditory cortex subareas vary substantially across individuals

The inherent significance of the functional variability of human auditory cortex has received relatively little attention. Here, we showed that fine-grained functional regions of auditory areas in STC vary substantially between individuals (Dice for resting state data: 57–68%). Notably, the parcellations were highly reproducible within individuals (Dice for resting state data: 69–78%), which indicates that the parcellations were reliable and captured idiosyncratic features that are related to brain function rather than noise. Functional homogeneity analysis of the resting-state data provided further evidence for neurophysiological validity by showing that the individual-specific parcellation results in functionally more homogeneous networks than the group-level parcellation or the individual-level parcellations not specific to the participant of interest. Additionally, the individual-specific parcellation yielded more precise correspondence within each individual’s task response topography than with group level parcellation or the parcellations of the other participants. This suggests that individual variability in parcellations corresponds with individual variability in task-evoked functional representations and, therefore, in the function of the auditory cortex. Our results are consistent with studies that have shown correspondences between task-evoked fMRI-response patterns and individual-specific whole-brain functional network parcellations [48, 84–86]. The individual variability found in auditory cortex parcellations is also in line with a recent study that found intrinsic functional connectivity patterns of the human auditory cortex to be highly variable across individuals [30].

The interindividual variability of the auditory cortex parcellations was larger in the left (Dice for resting-state data: 57–62%) than in the right hemisphere (Dice for resting-state data: 62–68%). This lateralization pattern is consistent with a recent fMRI study comparing individual variability of functional connectivity in humans and macaques [30], as well as with earlier studies reporting high interindividual variability in the left STG activity during word repetition task [87]. One potential interpretation is that the left auditory cortex is more individually variable due to its contribution to language related functions, which are shaped by our life-long experiences and exposure to language.

Traditional volume-based registration methods rely on landmarks defined by three-dimensional (3D) macroanatomy of structural MRI. However, the 3D macroanatomical shape of auditory areas differs greatly between individuals, which could confound estimates of interindividual functional variability. For example, Heschl’s gyrus may consist of one to three distinct transversal folds [29, 88, 89]. Therefore, in this study, the data were aligned between participants in the surface space, based anatomical registration methods that are adapted to each subject’s individual folding patterns [90]. This surface-based method has shown to largely remove macro-anatomical variability and result in highly consistent anatomical representations across participants [91, 92]. Thus, it has been proposed that the possible remaining variability in functionally defined areas across participants should reflect actual functional variability [92]. In line with these studies, our results show that the dissimilarity in the parcellations was not explained by the dissimilarity of macroanatomical properties (p>0.34). The individual-level functional connectivity-based mapping also provides a complementary method for aligning data across participants according to their functional instead of anatomical landmarks.

### 3.4 Auditory cortex functional connectivity networks are shaped by tasks but remain consistent within individuals

The reproducibility between the rest- and task-based parcellations (Dice: 62–74%) was lower than the reproducibility between rest-based parcellations (Dice: 69–78%), suggesting that the task performance can shape the functional connections between brain areas. This result is not fully consistent with the previous study reporting no differences between task and resting-state whole-cortex parcellations [51]. One reason for the contradicting results may be that we used more fine-grained parcellations and higher resolution 7T fMRI data, allowing the detection of more subtle differences. However, the parcellations derived from task data were more similar with the parcellations generated from the same participant’s resting-state data than with the task-based parcellations of the other participants. Thus, the functional organization of the brain is still most consistent within individuals regardless of how the brain is engaged in a task during scanning. This finding is consistent with a previous study showing that individuals can be identified from a large group based on their functional connectivity profiles regardless of the task conditions [93].

### 3.5 Limitations and future directions

Several limitations should be noted when interpreting the results of this study. First, the number of networks was selected by relying on technical criteria and may not be physiologically meaningful for each individual. The number of networks can differ between individuals particularly when studying patients who have undergone functional reorganizations. To overcome this issue, the parcellation algorithm could be initialized from a group-level atlas with a large number of networks (e.g., 40), followed by a gradual refinement of the number of subdivisions by combining regions with similar time courses (e.g., Pearson correlation r > 0.5). Second, the functional parcellation was based solely on STC-cortical connectivity. Given the converging evidence of the participation of subcortical structures [94–96] and cerebellum [97, 98] in auditory processing, future work should incorporate STC-subcortical and STC-cerebellar connectivity into the parcellation. Third, the parcellations should still be validated with other modalities, such as electrophysiological measurements or cortical stimulation. Fourth, to understand the functional meaning of the networks, the parcellations outlines should be compared with task activation topographies of more comprehensive set of auditory stimuli. Finally, our results indicated high interindividual variability in the functional organization of auditory cortex and showed that this variability reflects variability in the processing of auditory information in the brain. To understand more precisely how this interindividual variability is reflected in auditory function, hearing abilities, and speech comprehension, the results should be compared with behavioral data from hearing and speech comprehension tests.

## 4 Conclusion

Our results describe individual-level functional connectivity-based parcellations of human auditory areas in STC at three different network resolutions. These parcellations were highly reproducible within participants and variable between participants, suggesting that the parcellations are reliable and show meaningful interindividual variability. Furthermore, the parcellation specific to the individual yielded higher alignment with task response topographies of the same individual than with the parcellations of the other participants or group-level parcellation. This suggests that individual variability in parcellations reflects individual variability in auditory function. Our results extend previous literature of auditory cortex parcellation which have mainly focused on tonotopic mapping or group-level parcellations, derived from anatomical data. The high interindividual variability of the parcellations suggests that novel strategies considering the individuality of functional arrangements of auditory cortex could provide a way to facilitate future studies of human auditory cognition.

## 5 Materials and Methods

### 5.1 Participants

Thirty healthy volunteers (32.4 ± 10 years, 15 women, all right-handed) were recruited in this study via the Rally website that advertises internal clinical research studies of Massachusetts General Brigham (MGB). Twenty-eight of the participants were native English-speakers. None of the participants reported the use of cognition-altering medications, difficulties in hearing, or exposure to excessive noise. The study protocol was approved by the Institutional Review Board at MGB. The study was carried out in accordance with the guidelines of the declaration of Helsinki. A written informed consent was obtained from all participants prior to participation. Twenty of the participants participated in an additional hearing test. Based on Hughson-Westlake pure tone procedure, all but one participant had a hearing threshold of under 25 dB at 0.125–8 kHz. One of the participants had a 40 dB threshold at the highest frequencies of 6 and 8 kHz in the left ear and 35 dB at the frequency of 8 kHz in the right ear. Because of our goal to develop and test an individualized functional parcellation strategy for wide purposes, this participant was included in the study. Further, similarly to all other participants, this participant performed within a normal range in a computerized, automatic speech-in-noise comprehension task (QuickSIN test procedure; SNR loss: mean: –0.44, standard error of the mean: 0.2, range: –1.8–0.9). The supplemental hearing tests were conducted using a Diagnostic Audiometer AD 629 (Interacoustics, Middelfart, Denmark) with IP30 insert earphones.

### 5.2 Experimental protocol

Twenty-two participants completed three fMRI sessions on different days, and eight completed them in four days. Two of the sessions included six 8-minute resting-state fMRI scans (for one participant, only four scans were performed in one resting-state session). The additional tasks which were performed in up to two separate sessions, included a combined tonotopy and amplitude modulation (AM) rate representation task (2 × 8 min) and an audiovisual speech perception task (4 × 11 min). The duration of each session, with preparation, was around two hours. The participants were instructed to keep their eyes open and avoid any movement during all scans. During the resting-state scans, the participants were presented with a white fixation cross on a black background. In the task sessions, participants were instructed to attentively perform the given auditory task by indicating their responses through the use of an MRI-compatible response pad.

In the "Tonotopy/AM task," the participants were presented with serial blocks consisting of seven sounds, which varied both in their center frequency (tonotopy mapping) and in their AM rate (rate representation mapping). The carrier sounds were octave-wide bands of filtered white noise whose center frequencies were 0.12, 0.47, 1.87, or 7.47 kHz. These four possible carrier signals were modulated at rates centered either at around 4 Hz (*slow AM*) or at around 32 Hz (*fast AM*). In 85% of these blocks, which were non-targets, the AM rate was varied randomly at seven possible 1/8 octave steps around, and including, the center of the AM rate window, akin to an "AM melody" (adapted and modified from [99]. In the remaining 15%, which were the target blocks, the AM rate was constantly either 4 or 32 Hz. Participants were asked to look at a fixation mark at the screen, ignore the changes in the carrier frequency (i.e., the tonotopy aspect), pay attention to the AM rate of the sounds, and press a button with their right index finger upon hearing a target block with a constant AM rate. Each 8-minute run included 64 sound blocks, interleaved with silent baseline blocks.

In the "Audiovisual Speech/Noise task," the participants were presented with 636 (159 in each run) audiovisual recordings of a female talker speaking the words "rain" or "rock" [82, 83]. The auditory component was either acoustically intact or replaced with noise that matched the spectrotemporal power distribution of the original speech (for details of the stimulus creation, see [82]. Similarly, the visual component was either intact or a blurred counterpart of the original version. The two types of auditory and visual components were combined, resulting in four conditions for both "rain" and "rock": 1) Auditory Clear/Visual Clear, 2) Auditory Clear/ Visual Noisy, 3) Auditory Noisy/Visual Clear, and 4) Auditory Noisy/Visual Noisy. The stimuli were downloaded from https://openwetware.org/wiki/Beauchamp:Stimuli#Stimuli_from_Ozker_et_al. Participants were asked to press a button with their right index finger when hearing "rain" and another button with their right middle finger upon hearing "rock.”

The stimulus presentation was controlled with Presentation software (Neurobehavioral Systems, Berkeley, CA, USA). The auditory stimuli were delivered to the participants’ ears through MR-compatible insertable earphones (Sensimetrics Corporation, Model S15, Woburn, MA, USA). The intensity level of the stimuli was adjusted individually at a comfortable listening level that was clearly audible above the scanner noise. The fixation cross and the visual stimuli were projected to a mirror mounted on the head coil.

### 5.3 Data acquisition

The brain imaging data were acquired using a 7T whole body MRI scanner (MAGNETOM Terra, Siemens, Erlangen, Germany) and a home-built 64-channel receive brain array coil and a single-channel birdcage transmit coil [100]. To minimize head motion, participants’ heads were stabilized with MRI-compatible foam pads. In each scanning session, T1-weighted anatomical images were collected using a 0.75-mm isotropic Multi-Echo MPRAGE pulse sequence [101]; repetition time, TR = 2530 ms; echo time, 4 echoes with TEs 1.72, 3.53, 5.34 and 7.15 ms; flip angle = 7°; field of view, FoV = 240 × 240 mm^2^; 224 sagittal slices). To improve the accuracy of pial surface reconstruction, T2-weighted anatomical images were additionally acquired for 28 out of 30 participants with the T2 SPACE sequence (voxel size = 0.83 × 0.83 × 0.80 mm, TR = 9000 ms, TE = 269 ms, flip angle = 120°, FoV = 225 × 225 mm^2^, 270 sagittal slices) in one session.

Blood-oxygenation-level-dependent (BOLD) functional MRI data were collected using a single-shot gradient-echo blipped-CAIPI simultaneous multi-slice (SMS, acceleration factor in slice-encoding direction: 3, acceleration factor in phase-encoding direction: 4) 2D echo planar imaging (EPI) sequence [102]; TR = 2800 ms; TE = 27.0 ms; isotropic 1-mm^3^ voxels; flip angle = 78°; FoV = 192 × 192 mm^2^; 132 axial slices; phase enc. dir.: anterior→posterior; readout bandwidth = 1446 Hz/pixel; nominal echo spacing = 0.82 ms; fat suppression). For de-warping of the functional data, each session included an EPI scan collected with the same parameters, but in an opposite phase encoding polarity (posterior→anterior, PA-EPI) and a gradient-echo B0 field map (TR: 1040 ms, TEs: 4.71 ms and 5.73 ms; isotropic 1.3-mm^3^ voxels; flip angle: 75°; FoV: 240 × 240 mm^2^; 120 slices; bandwidth = 303 Hz/pixel).

During the fMRI data acquisition, participants’ heart rate and respiration signals were recorded at 400 samples/s. The heart rate was measured using the built-in Siemens photoplethysmogram transducers placed on the palmar surface of the participant’s index finger. Respiratory movements were recorded with the built-in Siemens respiratory-effort transducer attached to a respiratory belt, which was placed around the participant’s chest.

### 5.4 Data preprocessing

The bias field of the anatomical T1 and T2 images was first corrected using SPM12 (http://www.fil.ion.ucl.ac.uk/spm/, spm_preproc_run.m; bias full-width at half-maximum, FWHM: 18 mm, sampling distance: 2 mm, bias regularization: 1e-4) and custom MATLAB scripts. Thereafter, the cortical reconstructions were generated for each participant using recon-all function of FreeSurfer 7.1 [103] with an extension for submillimeter 7T data [104]. Both the T2 image and an average over 3–4 T1 images were used in the reconstruction to improve the quality of the cortical surfaces. For intracortical smoothing, nine additional surfaces were generated at fixed relative distances between the white and pial surfaces produced by recon-all. Finally, the quality of the reconstructed surface was reviewed for quality assurance, and pial and white mater segmentation errors were corrected using the Recon Edit function of Freeview.

fMRI analysis was conducted using FreeSurfer 7.1. The fMRI data were first slice-time and motion corrected. Geometric distortions induced by inhomogeneities in the static magnetic field were compensated by dewarping. The distortion field for dewarping was estimated based on the fMRI data and the PA-EPI scan which have equal and opposite distortions compared to the AP-EPI scans used for BOLD fMRI acquisition. From these scans, the susceptibility-induced off-resonance field offsets were estimated using a method described in [105] as implemented in the FMRIB Software Library (FSL) [106]; topup, applytopup, FSL 5.0.7). For four participants with a missing PA-EPI scan, the distortion field was estimated using the B0 field map (epidewarp, FreeSurfer 7.1). Heart rate and respiratory artifacts were corrected with RETROspective Image CORrection (RETROICOR) algorithm [107]; 3^rd^ order heart rate, respiratory, and multiplicative terms). For the three participants with missing heart rate recordings, only respiratory data were used in RETROICOR. RETROICOR was not applied on five participants’ data because of missing heart rate and respiration recordings. Boundary-based registration [108]; preproc-sess, FreeSurfer 7.1) was used to co-register the functional data with the anatomical images. The fMRI data were smoothed spatially using an intracortical smoothing procedure applied within the cortical gray matter to reduce noise contamination and signal dilution from surrounding cerebrospinal fluid and white matter, resulting in higher gray matter specificity compared to conventional three-dimensional volume-based smoothing [109–112]. For smoothing, the fMRI time series were resampled onto the 11 intermediate intracortical surfaces by projecting each intersecting voxel onto the corresponding surface vertices using trilinear interpolation. These surfaces were used as an input for the smoothing algorithm that generated one intracortically smoothed surface. The data were smoothed using a three-dimensional kernel that smooths tangentially with the 7^th^ order vertex neighborhood, and radially across 10 cortical surfaces. The top surface of the gray matter was excluded to reduce partial volume effects from the surrounding cerebrospinal fluid. Thereafter, the data were bandpass filtered between 0.01–0.08 Hz and signals from white matter, cerebrospinal fluid, and the area outside of the head were regressed out. The last two steps were performed only for the parcellation but not for GLM analysis since they may decrease the effect of interest.

### 5.5 Data analysis

#### 5.5.1 Group-based functional atlas

Group-level functional network atlases for the auditory areas were estimated using the same approach as in [61]. The parcellations were generated in the FreeSurfer average subject space (fsaverage6, 40,962 vertices per hemisphere). The fMRI time-series data from each individual subject was projected into this surface space, creating time-series data with 81,924 vertices. The STC area was defined by combining superior temporal and transverse temporal labels from Desikan-Killiany atlas [113]. For each STC vertex of every participant, its functional connectivity profile was estimated by computing the Pearson correlation between its time series and that of each of the other cortical vertices (including both hemispheres), i.e., each functional connectivity profile was an 81,923×1 vector of correlation coefficient values. Thereafter, the functional connectivity profiles of each STC vertex were averaged across participants, and a k-means clustering algorithm was applied to cluster the vertices into networks based on the cosine similarity of their functional connectivity profiles. To avoid getting trapped in local minima, the clustering was performed 500 times with randomly selected initial network centroids, and the clustering solution was selected that yielded the smallest aggregate distance. We varied the network number from 2 to 24 and selected for individual-level investigation four solutions that yielded local maxima of the product of their Dice and Silhouette coefficients (4-, 6, 11- and 19-network parcellations; Fig. S1 in Supplementary Material; 19-network parcellation was excluded from the main analyses since it was not reproducible at the individual level). To compute the Dice values for this analysis, the fMRI data from each run were split in half, and the first halves were concatenated together, and the last halves were concatenated together. Thereafter, the Dice values were computed between the two parcellations generated from these concatenated data. The Dice coefficient represents the percentage of vertices that were assigned into the same network in separate parcellations. The Dice coefficient was computed for each network in the parcellation as follows:

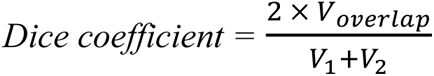

where 𝑉*_overlap_* is the number of vertices assigned into the same network in the parcellations, and 𝑉_1_ and 𝑉_2_ are numbers of vertices of the network in each of the parcellations. Silhouette value was used to measure how similar the functional connectivity profile of a vertex is to the other connectivity profiles in the same cluster compared to the connectivity profiles of the other clusters. The Silhouette coefficient was calculated as follows:

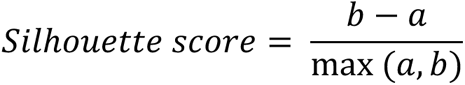

where 𝑎 is an average distance between each data point within a cluster and 𝑏 is the average distance between all clusters.

Clusters of size 10 mm^2^ or smaller were reassigned to larger clusters. To this end, the network assignment was computed for six neighboring vertices for each vertex within these small clusters. Thereafter, each vertex was assigned to the network that was the most frequent among the neighboring vertices. If there were equally frequent networks among the neighbors, a Pearson correlation was computed between the functional connectivity profile of the vertex and the average functional connectivity profile of the most frequent networks. The vertex was assigned to the network with which the correlation was highest. This procedure resulted in reassignment of one vertex in the left hemispheric 4-network parcellation and 5 vertices in the left hemispheric 11-network parcellation.

#### 5.5.2 Individual-level parcellation

Individual-level parcellations were generated by adjusting the functional connectivity-based group parcellation using the same procedure as [51]. 1) Individual participant’s fMRI data was projected into the fsaverage6 surface (FreeSurfer, mri_surf2surf), and fMRI signal time courses were averaged across the vertices that fell within each network of the group-level functional network parcellation. These network time courses were used as the “reference signals” for the following individual-level adjustment procedure. 2) Each vertex of an individual participant was reassigned to one of the networks according to its maximal correlation with the corresponding reference signals. A confidence level was computed for each reassignment by dividing the strongest correlation value by the second strongest one. 3) A ‘high-confidence signal’ was created for each network by averaging all vertices that were reassigned to that network that exceeded a confidence value of 1.3. 4) The high-confidence and original reference signals derived from the group parcellation were averaged, and the resulting functional connectivity profiles were used as new reference signals for the next iteration. This allowed us to use both individual participant’s information and the information of the group parcellation to guide the reassignment of the vertices. Finally, steps 3–4 were iterated ten times, and the end result was used as the individual parcellation. Ten iterations were used since the parcellations start to became stable approximately at the 5th iteration (dice values of resting state parcellations between 9^th^ and 10^th^ iterations: left: 4 networks: 99±1%, 6 networks: 99±1%, 11 networks: 97±3%; right: 4 networks: 99±0%, 6 networks: 99±1%, 11 networks: 97±4%), and high number of iterations increases the risk of overfitting the data. Individual-level parcellations were generated for 4-, 6-, and 11-network group parcellations using the fMRI data from 1) the first restring-state session, 2) second resting-state session, 3) task session, and 4) all sessions. The reproducibility of the parcellations was estimated by comparing the parcellations generated based on different sessions.

We rejected one network from the left and one from the right hemisphere of the individual-level 11-network parcellations since they were not reproducible. Moreover, they did not appear to be neurophysiologically plausible since they included clusters of very few vertices distributed across the STS. The corresponding contralateral networks were reproducible and, therefore, they were not rejected. These networks were also unilateral in the 11-network group parcellation. The other networks were bilateral both in the individual and group parcellations.

#### 5.5.3 Intraparticipant reproducibility and interindividual variability of the parcellations

Intraparticipant reproducibility and interindividual variability of the parcellations were estimated with the Dice coefficient. The Dice coefficient for the parcellation as a whole was computed by averaging the network-specific Dice coefficients. To estimate the reproducibility of the parcellations, the Dice coefficients were computed between the parcellations derived from the two resting-state sessions collected on different days. We also tested whether the parcellation algorithm can generalize to different tasks by computing Dice coefficients between the parcellations generated based on resting-state and task fMRI data. The interindividual variability was determined by averaging the Dice coefficients computed between each participant and all other participants within the resting-state sessions. Further, we compared the Dice coefficients between task and resting-state parcellations of the same participant with the Dice coefficients between task parcellations of the participant and any other participants. The differences were estimated with Friedman test (friedman, MATLAB) and pairwise Wilcoxon signed rank tests (signrank, MATLAB). The results were corrected for multiple comparisons with Benjamini-Hochberg procedure [55, 114]. Individual-level analysis revealed that many networks of the 19-network parcellation were not reproducible and, therefore, we excluded the 19-network parcellation from further analysis.

The Mantel test [115] was used to test whether the interindividual variability of the parcellations reflects variability in the cortical thickness or curvature. The thickness and curvature maps for each participant were created by the FreeSurfer recon-all pipeline. The similarity matrices were computed for the curvature and thickness by computing pairwise Pearson correlations between participants for the values within STC. The similarity matrix describing parcellation similarity was created from the Dice values of the parcellation created using all resting state and task sessions. The Spearman correlation was computed between the upper triangle entries of the Dice similarity and thickness similarity matrices as well as between the Dice similarity and curvature similarity matrices. The statistical significance of the results was estimated by recalculating the correlations for 10,000 random permutations of the rows and columns of the Dice similarity matrix with respect to another.

#### 5.5.4 Comparing parcellations with brain areas activated by auditory or audiovisual stimulation

GLM analysis was conducted with the fMRI analysis stream of Freesurfer, fs-fast, to estimate brain areas activated during auditory and audiovisual tasks. In the Tonotopy/AM task, the following contrasts were computed: 1) auditory stimulation vs. baseline, 2) high (1.87 kHz or 7.47 kHz) vs. low (0.12 kHz or 0.47 kHz) carrier frequencies, 3) slow (4 cycles/s) vs. fast (32 cycles/s) amplitude modulation, 4) slow amplitude modulation vs. baseline, and 5) fast amplitude modulation vs. baseline. The occasional target stimuli were excluded from all contrasts.

In the Audiovisual Speech/Noise task, we concentrated on three contrasts computed across the Auditory Clear / Visual Clear (AcVc), Auditory Clear/ Auditory Noisy (AcVn), Auditory Noisy/ Visual Clear (AnVc), and Auditory Noisy/Visual Noisy (AnVn) stimuli. In main effect of Speech vs. Noise, we contrasted all possible audiovisual combinations with clear auditory signal with those with noisy auditory signal, i.e., ((AcVc+AcVn)-(AnVc+AnVn))/2. In the main effect of Lip motion vs. Noise, all audiovisual combinations with clear visual signal were contrasted with those with blurred vision, i.e., ((AcVc+AnVc)-(AcVn+AnVn))/2 We also calculated an audiovisual interaction that was specifically aimed at identifying the cortical areas where visual information of lip motion has the strongest effect of processing of speech sounds when the auditory signal is noisy. This contrast was, thus, defined as (AnVc-AnVn) - (AcVc-AcVn).

The group-level results were computed with a weighted least-squared random-effects model. If the task data were measured from the participant on two different days (Tonotopy/AM task for three participants, Audiovisual Speech/Noise task for seven participants), the measurements were combined with a fixed-effect model before computing the group-level results. The results were corrected for multiple comparisons with a cluster-wise Monte Carlo simulation (cluster-wise *p*-value threshold: *p*<0.05, voxel-vise threshold: 1.3, sign: absolute, 10,000 simulations).

The outlines of the parcellations were overlaid on GLM task contrast maps at individual and group levels to visually estimate whether the parcellations align with the activated brain areas. Functional inhomogeneity was further used to evaluate the inhomogeneity of task activation in each network [85]. Functional inhomogeneity of the parcellation was estimated by calculating the standard deviation in fMRI task activation (*z*-scores) within each network and averaging the standard deviations across networks while accounting for the network size:

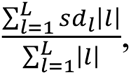

where 𝑠𝑑*_l_* is the standard deviation of task activation *z*-values for network *l* and |𝑙| is the number of vertices in the network *l* [85].

The functional inhomogeneity was computed for each participant, using their task activation map with 1) their own individual-specific parcellation, 2) the group average parcellation, and 3) the individual-specific parcellations of all other participants. The functional inhomogeneities between participants’ task activation and other participants’ parcellations were averaged across participants. The Friedman test (friedman, MATLAB) and pairwise Wilcoxon signed rank tests (signrank, MATLAB) were used to estimate differences in the functional inhomogeneity between the three parcellations. The results were corrected for multiple comparisons with Benjamini-Hochberg procedure [55, 114].

Functional specificity of the STC parcellations was additionally evaluated for each task contrast by calculating the percentage of the total task effect each of the networks explained in each parcellations. This was done by dividing the sum of the GLM contrast effect size within each network by the sum of the GLM contrast effect size values within the whole parcellation. We also calculated Pearson correlation of these network-specific percentages of the total contrast effect size between tasks. This allowed us to evaluate whether the networks explaining most of the task effect differ between tasks.

### 5.6 Connectional homogeneity

To compute the connectional homogeneity for the parcellations, 12 resting-state fMRI runs were first concatenated within each participant. Second, pairwise Pearson correlation coefficients were computed between all vertices within each network of every participant. Third, these pairwise correlation coefficients were transformed into z-values using Fisher transformation. Finally, the z-values were averaged within each network, and thereafter, the averaged z-values were further averaged across all networks while accounting for network size:

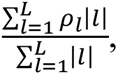

where 𝜌*_l_* is the connectional homogeneity for network *l* and |𝑙| is the number of vertices in the network *l* [85].

The connectional homogeneity was computed for each participant, using their resting-state data and 1) their own individual-specific parcellation, 2) the group average parcellation, and 3) the individual-specific parcellations of all other participants. The functional connectivity homogeneities between each participant’s data and other participants’ parcellations were averaged. The Friedman test (friedman, MATLAB) and pairwise Wilcoxon signed rank tests (signrank, MATLAB) were used to determine whether the connectional homogeneity within networks differed between parcellations. The results were corrected for multiple comparisons with Benjamini-Hochberg procedure [55, 114].

### 5.7 Seed-based functional connectivity

Seed-based functional connectivity analysis was conducted to estimate with which cortical areas STC networks were connected. A seed-based connectivity map was generated for each network using that network as a seed region. To this end, the fMRI signals were averaged across vertices within the seed network as the seed signal. The functional connectivity map for the seed region was estimated in fsaverage6 surface by computing Pearson’s correlation between the seed signal and the fMRI signal from each cerebral cortical vertex of the same hemisphere. For the statistical analysis, the correlation values were normalized using Fisher *z*-transformation [116]. The statistical significance of the connectivity maps was estimated with one-sample t-test (FreeSurfer, mri_glmfit). The statistical maps were corrected for multiple comparisons with the cluster extent correction using 5000 permutations and a voxel-wise cluster-forming threshold of 8. Further, we estimated pairwise differences between seed-based functional connectivity maps for adjacent networks in the 11-network parcellation using the Wilcoxon signed-rank test and Benjamini-Hochberg correction for multiple comparisons [55].

## Acknowledgments

Our work was funded by NIH grants R011DC016915, R01DC016765, R01DC017991, S10OD023637, R01DC016765, R01DC017915, R01DC017991, P41EB015896, CIHR MFE-171291, Finnish Cultural Foundation, Changping Laboratory (2021B-01-01) and China Postdoctoral Science Foundation (2022M720529 and 2023M730175). This work was additionally supported in part by the NIH NIBIB (grant P41-EB030006) and by the MGH/HST Athinoula A. Martinos Center for Biomedical Imaging; and was made possible by the resources provided by NIH Shared Instrumentation Grants S10-OD023637. Much of the computation resources required for this research was performed on computational hardware generously provided by the Massachusetts Life Sciences Center (https://www.masslifesciences.com/).

