## Supplementary material for "Individual connectivity-based parcellations reflect functional properties of human auditory cortex"

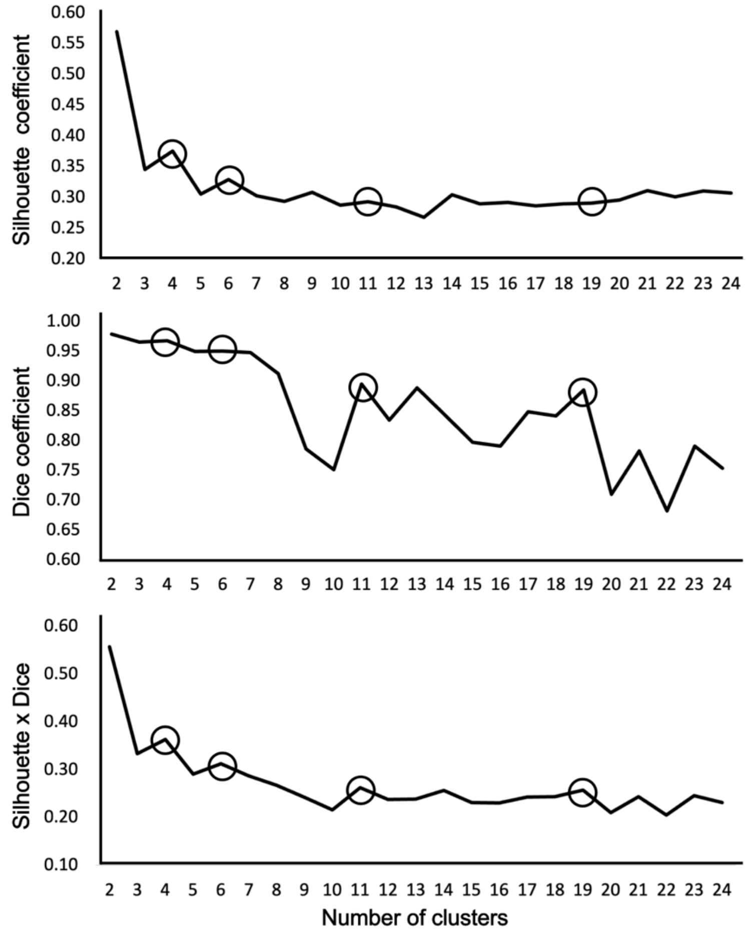


***Figure S1.*** *Silhouette and Dice coefficients as well as their product computed for parcellations with parcel numbers from 2 to 24. Higher Silhouette coefficient means higher separability between clusters and higher Dice coefficient higher reproducibility of the parcellations within participant. The parcellations selected for further investigation are marked with circles.*

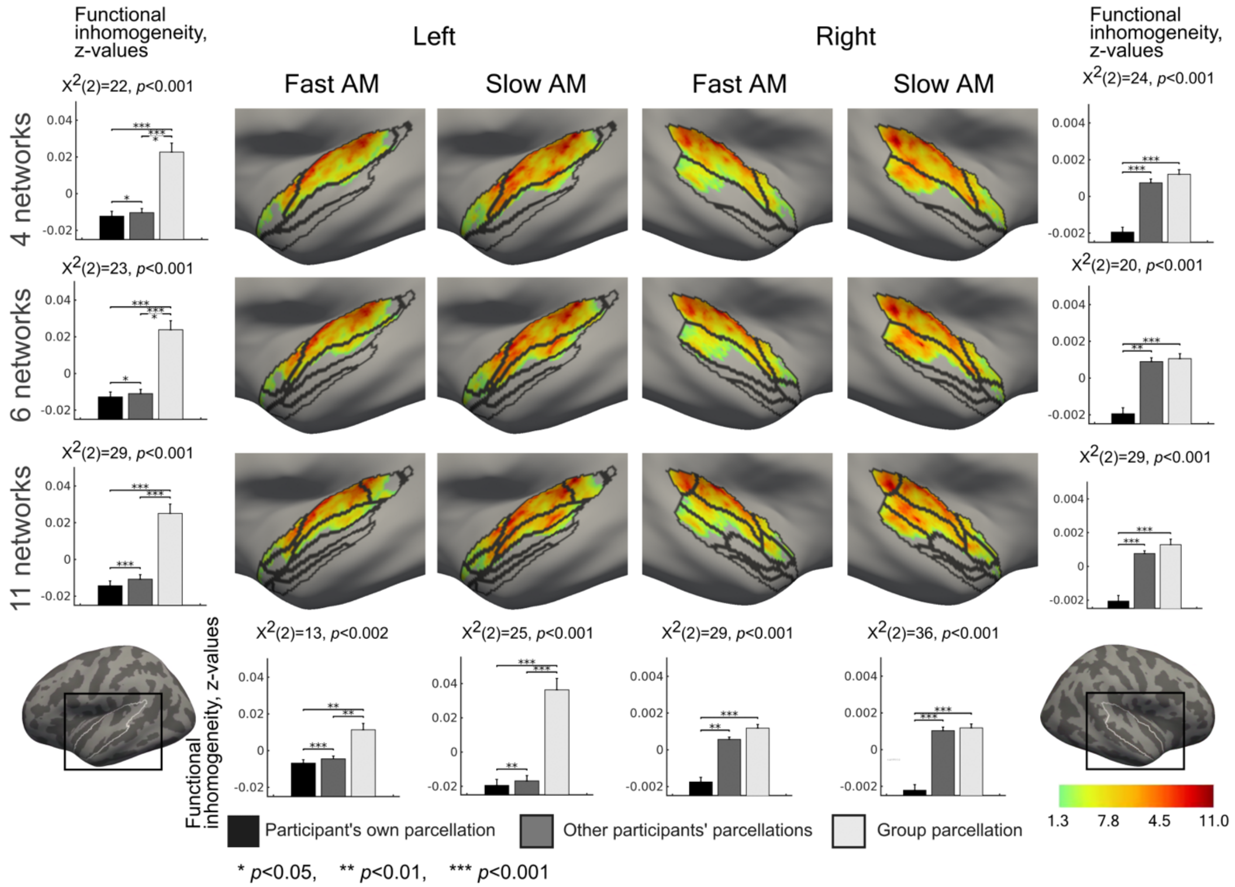


***Figure S2.*** *Topographic correspondence between the STC parcellations and amplitude modulation rates, overlaid with the three studied group-level parcellations. The strongest activation to fast AM appears to concentrate to the more medial aspect of early auditory cortex than those for the slow AM. The bar diagrams in the right and left columns show right- and left-hemispheric functional inhomogeneity averaged over fast and slow AM contrast maps within each of the three parcellations. The bar diagrams in the bottom row show functional inhomogeneity averaged over all parcellations within each of the four contrasts. The functional inhomogeneity was computed between the individual GLM contrast map of each participant and 1) their own individual–specific parcellation, 2) individual-specific parcellation of all other participants, and 3) group-average parcellation. The differences between these three conditions were estimated with the Friedman test and pairwise Wilcoxon signed rank tests. The results were corrected for multiple comparisons using Benjamini-Hochberg procedure. Error bars indicate standard errors of the mean (SEM).*

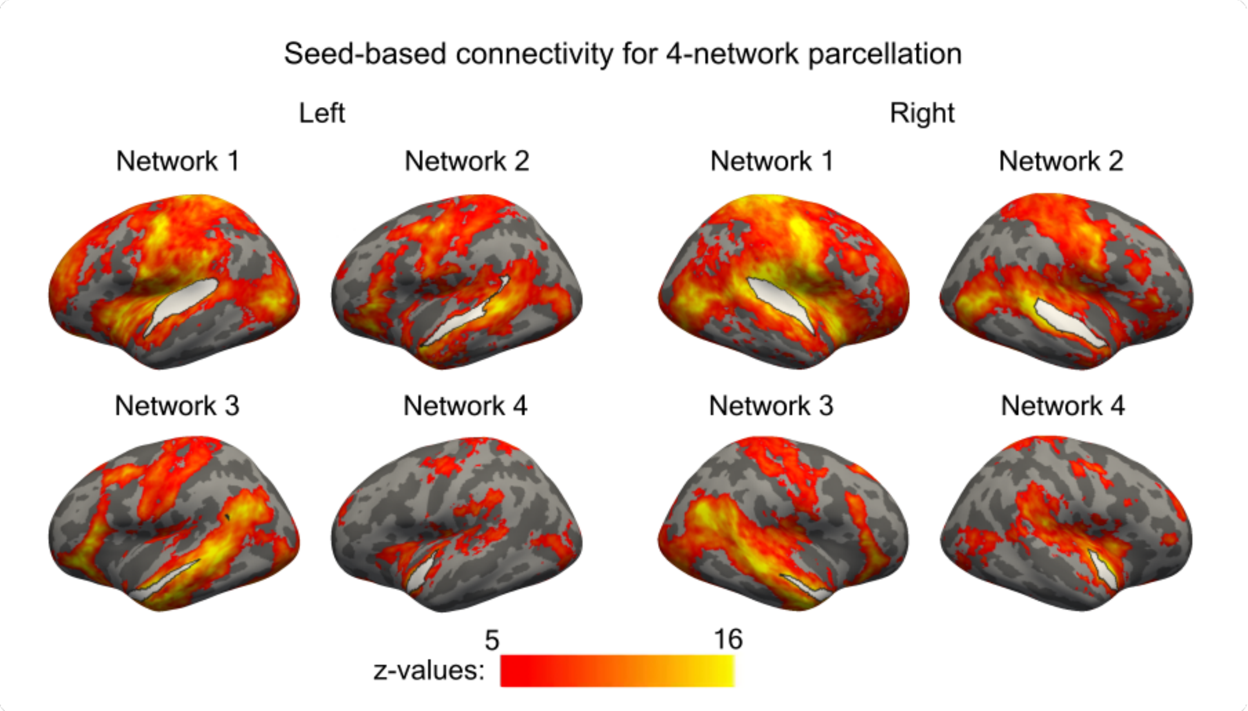


***Figure S3.*** *Seed-based functional connectivity maps for each parcel in 4-parcel parcellation One-sample t-test was performed for each vertex. The presented maps are thresholded at p<0.05 and corrected for multiple comparisons with cluster-extent based permutation thresholding with a cluster-forming threshold of p<0.001 (one-sample T-test).*

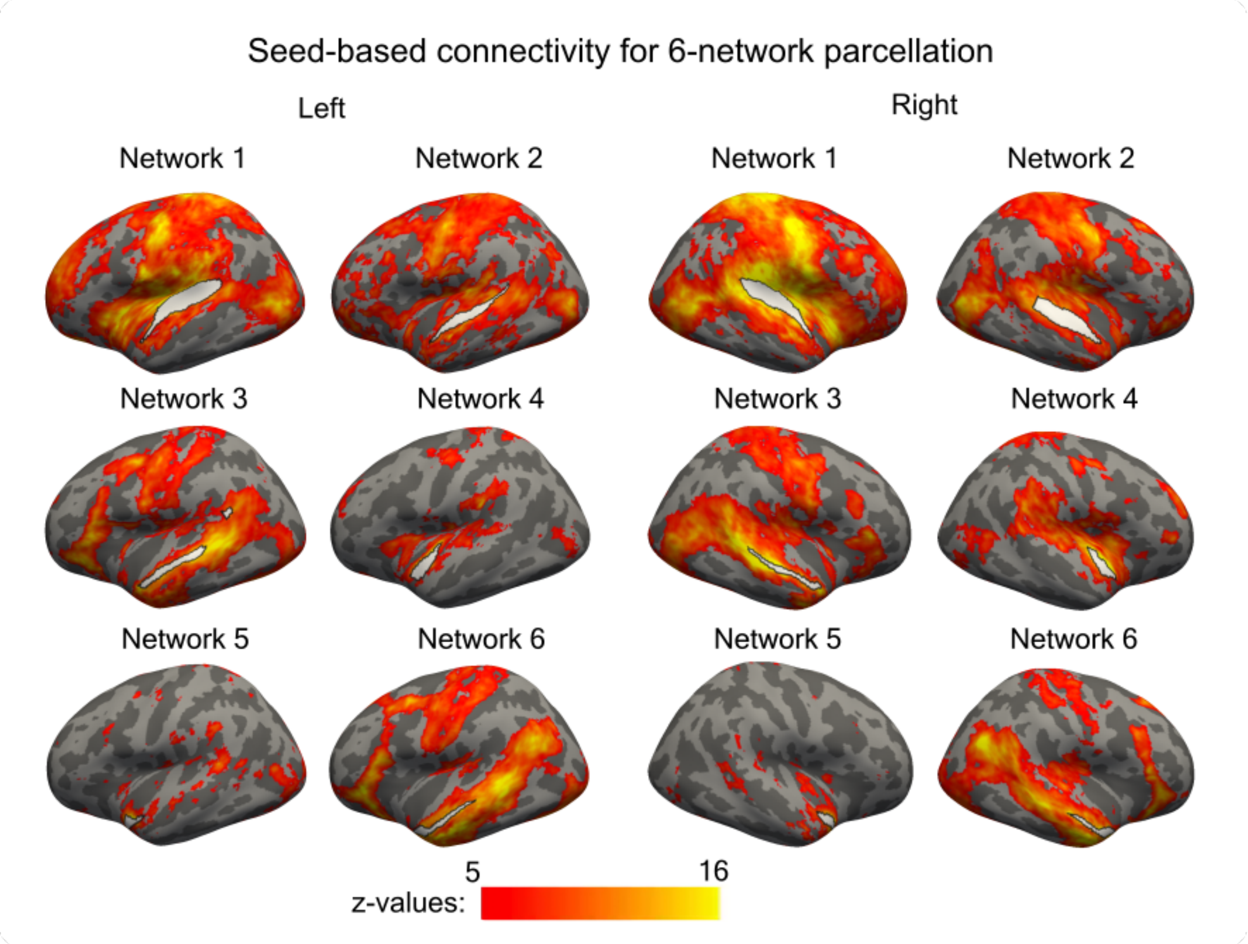


***Figure S4.*** *Seed-based functional connectivity maps for each parcel in 6-parcel parcellation. One-sample t-test was performed for each vertex. The presented maps are thresholded at $p<0.05$ and corrected for multiple comparisons with cluster-extent based permutation thresholding with a cluster-forming threshold of $p<0.001$ (one-sample T-test).*

***Table S1.*** *Dice coefficients (mean ± standard error of the mean, SEM) for resting-state parcellations within and between individuals.*

| Number of networks | Hemisphere | Resting state, intraindividual (%) | Resting state, interindividual (%) | *p*-value |
|---|---|---|---|---|
| 4 | Left | 74±1.4 | 62±0.8 | <0.001 |
| 6 | Left | 69±1.2 | 57±0.6 | <0.001 |
| 11 | Left | 69±1.0 | 57±0.5 | <0.001 |
| 4 | Right | 78±1.1 | 68±0.6 | <0.001 |
| 6 | Right | 76±1.1 | 64±0.6 | <0.001 |
| 11 | Right | 72±0.9 | 62±0.6 | <0.001 |

***Table S2.*** *Dice coefficients (mean ± SEM) between resting-state and task parcellations within participants and task parcellations between participants.*

| Number of networks | Hemisphere | Task vs. rest, intraindividual (%) | Task, interindividual (%) | p-value |
|---|---|---|---|---|
| 4 | Left | 67±1.4 | 61±1.0 | <0.002 |
| 6 | Left | 65±1.1 | 58±1.0 | <0.001 |
| 11 | Left | 62±0.9 | 55±1.0 | <0.001 |
| 4 | Right | 74±1.0 | 66±0.8 | <0.001 |
| 6 | Right | 71±1.0 | 63±0.8 | <0.001 |
| 11 | Right | 68±0.8 | 60±0.6 | <0.001 |

***Table S3.*** *Dice coefficients (mean ± SEM) between the two resting-state parcellations as well as between task and resting-state parcellations within individuals.*

| Number of networks | Hemisphere | Resting state, intraindividual (%) | Task vs. rest intraindividual (%) | p-value |
|---|---|---|---|---|
| 4 | Left | 74±1.4 | 67±1.4 | <0.010 |
| 6 | Left | 69±1.2 | 65±1.1 | <0.099 |
| 11 | Left | 69±1.0 | 62±0.9 | <0.006 |
| 4 | Right | 78±1.1 | 74±1.0 | <0.043 |
| 6 | Right | 76±1.1 | 71±1.0 | <0.014 |
| 11 | Right | 72±0.9 | 68±0.8 | <0.015 |

***Table S4.*** *Dice coefficient (mean ± SEM) for each parcel of the 4-parcel parcellation. Intraindividual Dice coefficients were calculated between resting state parcellations created from the two resting state sessions of the same participant. Interindividual Dice coefficients were calculated between resting state-parcellations of different participants within the two sessions. Dice coefficients were averaged over the sessions. P-value is the significance of the difference between Dice values (Wilcoxon signed rank test, corrected for multiple comparisons using Benjamini-Hochberg procedure across all parcellations and networks).*

| Parcel | Hemisphere | Intraindividual (%) | Interindividual (%) | p-value |
|---|---|---|---|---|
| 1 | left | 70±0.6 | 52±0.2 | <0.001 |
| 2 | left | 77±0.4 | 64±0.2 | <0.001 |
| 3 | left | 70±0.5 | 61±0.2 | <0.001 |
| 4 | left | 79±0.4 | 68±0.2 | <0.001 |
| 1 | right | 74±0.5 | 60±0.2 | <0.001 |
| 2 | right | 78±0.3 | 70±0.1 | <0.001 |
| 3 | right | 78±0.4 | 68±0.2 | <0.001 |
| 4 | right | 84±0.3 | 74±0.2 | <0.001 |

***Table S5.*** *Dice coefficient (mean ± SEM) for each parcel of the 6-parcel parcellation. Intraindividual Dice coefficients were calculated between resting state parcellations created from the two resting state sessions of the same participant. Interindividual Dice coefficients were calculated between resting state-parcellations of different participants within the two sessions. Dice coefficients were averaged over the sessions. P-value is the significance of the difference between Dice values (Wilcoxon signed rank test, corrected for multiple comparisons using Benjamini-Hochberg procedure across all parcellations and networks).*

| Parcel | Hemisphere | Intraindividual (%) | Interindividual (%) | p-value |
|---|---|---|---|---|
| 1 | left | 69±0.4 | 56±0.2 | <0.001 |
| 2 | left | 68±0.4 | 60±0.2 | <0.001 |
| 3 | left | 65±0.7 | 47±0.2 | <0.001 |
| 4 | left | 74±0.4 | 66±0.2 | <0.001 |
| 5 | left | 76±0.5 | 65±0.2 | <0.001 |
| 6 | left | 63±0.6 | 49±0.2 | <0.001 |
| 1 | right | 78±0.4 | 66±0.2 | <0.001 |
| 2 | right | 77±0.4 | 69±0.2 | <0.001 |
| 3 | right | 69±0.5 | 52±0.2 | <0.001 |
| 4 | right | 76±0.5 | 64±0.2 | <0.001 |
| 5 | right | 83±0.3 | 73±0.1 | <0.001 |
| 6 | right | 73±0.5 | 58±0.2 | <0.001 |

***Table S6.*** *Dice coefficient (mean ± SEM) for each parcel of the 11-parcel parcellation. Intraindividual Dice coefficients were calculated between resting state parcellations created from the two resting state sessions of the same participant. Interindividual Dice coefficients were calculated between resting state-parcellations of different participants within the two sessions. Dice coefficients were averaged over the sessions. Parcel 4 was rejected from the left and Parcel 3 from the right hemisphere and, therefore, they are not shown in the table. P-value is the significance of the difference between Dice values (Wilcoxon signed rank test, corrected for multiple comparisons using Benjamini-Hochberg procedure across all parcellations and networks).*

| Parcels | Hemisphere | Intraindividual (%) | Interindividual (%) | p-value |
|---|---|---|---|---|
| 1 | left | 60±0.5 | 45±0.2 | <0.001 |
| 2 | left | 75±0.4 | 68±0.2 | <0.001 |
| 3 | left | 65±0.6 | 53±0.2 | <0.001 |
| 5 | left | 73±0.6 | 57±0.2 | <0.001 |
| 6 | left | 65±0.7 | 52±0.3 | <0.001 |
| 7 | left | 72±0.6 | 64±0.2 | <0.012 |
| 8 | left | 61±0.6 | 44±0.2 | <0.001 |
| 9 | left | 78±0.4 | 70±0.2 | <0.019 |
| 10 | left | 76±0.3 | 67±0.2 | <0.001 |
| 11 | left | 62±0.6 | 47±0.2 | <0.001 |
| 1 | right | 66±0.6 | 55±0.2 | <0.001 |
| 2 | right | 77±0.4 | 71±0.2 | <0.036 |
| 4 | right | 77±0.4 | 63±0.2 | <0.001 |
| 5 | right | 76±0.5 | 68±0.2 | <0.001 |
| 6 | right | 66±0.6 | 58±0.3 | <0.005 |
| 7 | right | 80±0.4 | 68±0.2 | <0.001 |
| 8 | right | 63±0.6 | 48±0.2 | <0.001 |
| 9 | right | 80±0.4 | 67±0.2 | <0.001 |
| 10 | right | 74±0.5 | 69±0.2 | <0.004 |
| 11 | right | 63±0.7 | 48±0.2 | <0.001 |
